# On the heterozygosity of an admixed population

**DOI:** 10.1101/820241

**Authors:** Simina M. Boca, Lucy Huang, Noah A. Rosenberg

## Abstract

A population is termed *admixed* if its members possess recent ancestry from two or more separate sources. As a result of the fusion of source populations with different genetic variants, admixed populations can exhibit high levels of genetic variation, reflecting contributions of their multiple ancestral groups. For a model of an admixed population derived from *K* source groups, we obtain a relationship between its level of genetic variation, as measured by heterozygosity, and its proportions of admixture from the various source populations. We show that the heterozygosity of the admixed population is at least as great as that of the least heterozygous source population, and that it potentially exceeds the heterozygosities of *all* of the source populations. The admixture proportions that maximize the heterozygosity possible for an admixed population formed from a specified set of source populations are also obtained under specific conditions. We examine the special case of *K* = 2 source populations in detail, characterizing the maximal admixture in terms of the heterozygosities of the two source populations and the value of *F*_*ST*_ between them. In this case, the heterozygosity of the admixed population exceeds the maximal heterozygosity of the source groups if the divergence between them, measured by *F*_*ST*_, is large enough, namely above a certain bound that is a function of the heterozygosities of the source groups. We present applications to simulated data as well as to data from human admixture scenarios, providing results useful for interpreting the properties of genetic variability in admixed populations.

## 1 Introduction

Admixed populations are populations that possess ancestry from multiple source groups. They result from the fusion of populations that have long been separated, in processes such as long-distance migration and hybrid-zone formation at population boundaries.

Several features of ancestry and allele frequencies are characteristic of admixed populations (Chakraborty, 1986; Long, 1991; Verdu & Rosenberg, 2011; Gravel, 2012). In an admixed population, the values of allele frequencies are typically intermediate between those of the various sources. Unlike in a mixture that pools individuals taken from separate populations, in an admixed population, alleles from different sources cooccur within individuals. The contributions from the source populations are each large enough that most members of an admixed population have ancestry in more than one source group.

In admixed populations, the history of mating among populations is recent enough that time has not yet eroded differences among admixed individuals in their relative proportions of ancestry. This feature of high levels of variability in admixture proportions has been central to studies of admixed populations. Investigations of such phenomena as the timing and contributions of the source populations (Verdu & Rosenberg, 2011; Gravel, 2012), the effect of admixture levels on assortative mating patterns (Risch *et al.*, 2009; Zou *et al.*, 2015), and the genetic basis of traits in admixed populations (Buerkle & Lexer, 2008; Zhu *et al.*, 2008) all make use of variation in levels of admixture levels across admixed individuals.

A second aspect of variability in admixed populations is potentially of interest: the variability of alleles as captured by genetic diversity measures. The effect of admixture in contributing to increased genetic diversity, however, is not simple. For example, in a study of the genetics of populations founded by relatively small groups, Mooney *et al.* (2018) examined genetic diversity in admixed and non-admixed populations, some of which were regarded as founder populations. Mooney *et al.* (2018) observed that genetic diversity was relatively high in multiple admixed populations of Latin America. This pattern was observed even for populations that, on the basis of small population size and past history of isolation, might have been expected to have relatively low levels of genetic diversity.

Here, to deepen understanding of the relationship between admixture and genetic variability, we focus in admixed populations on levels of genetic diversity computed from allele frequencies, rather than on variability among individuals in admixture proportions. For a model of an admixed population with *K* source groups, we derive a relationship between genetic diversity, as measured by heterozygosity, and proportions of admixture drawn from the various source populations. The model is the same model we have previously used to examine the genetic differentiation between admixed populations and their source groups, as measured by *F*_*ST*_ (Boca & Rosenberg, 2011). We show that for all values of the admixture contributions from the source populations, the heterozygosity of the admixed population is greater than or equal to the smallest of the source population heterozygosities. We further examine the maximal values of the heterozygosity of the admixed population over the space of possible admixture proportions. We consider in more detail special cases of the admixture model with *K* = 2 and *K* = 3 source populations, providing explicit results for *K* = 2 in terms of relatively few parameters. Finally, we use simulations and example analyses from admixed human populations to illustrate the mathematical results.

## 2 Notation and model

We consider a model with *K* source populations and an admixed population arising from these sources. A single locus is considered, with *J* distinct alleles. In Sections 2.1, 2.2, and 2.3, respectively, we define the expected heterozygosity, fixation index, and allele-frequency dot product statistics that we use in our analysis. In Section 2.4, we introduce the admixture model. Notation is summarized in Table 1.

**Table 1:**
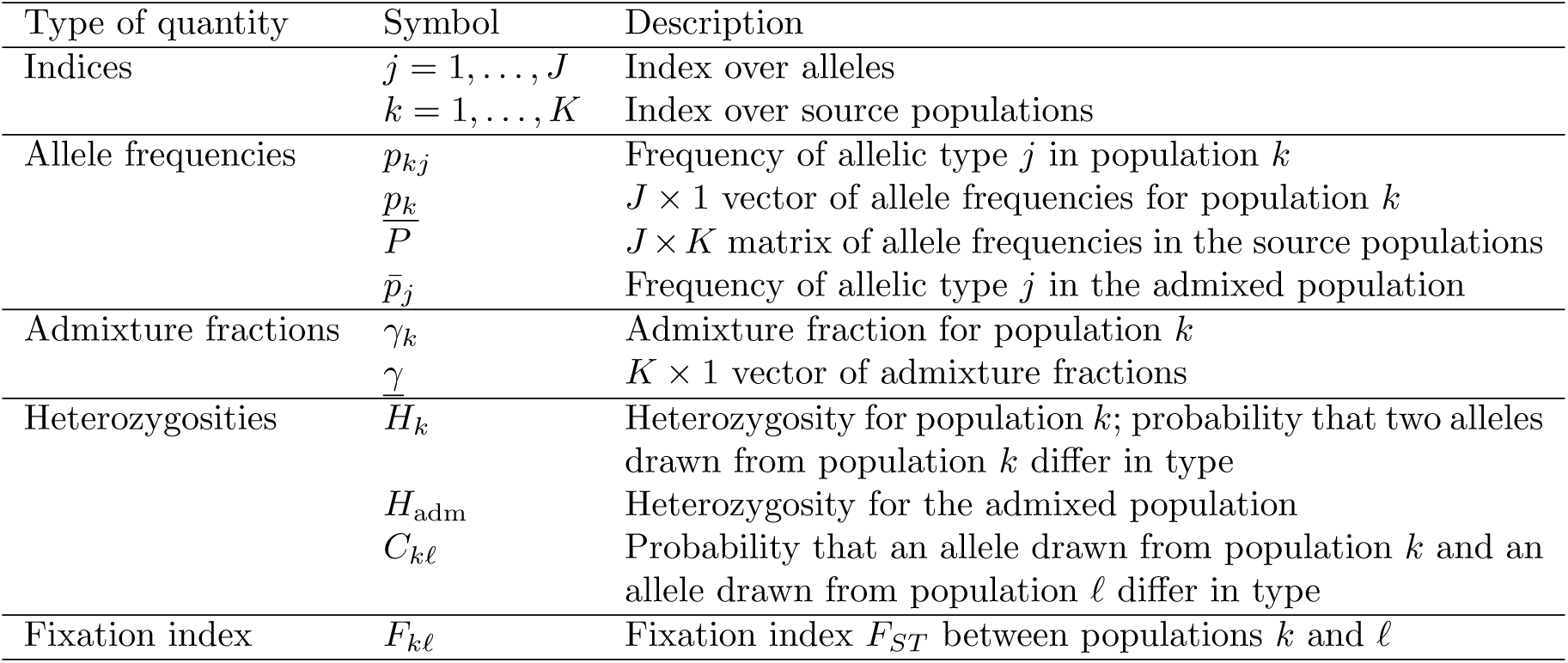
Notation.

### 2.1 Expected heterozygosity

We denote by *p*_*kj*_ the frequency of allelic type *j*, 1 ⩽ *j* ⩽ *J*, in source population *k*, 1 ⩽ *k* ⩽ *K*, with 0 ⩽ *p*_*kj*_ ⩽ 1. The expected heterozygosity is a measure of genetic diversity, giving the probability that two alleles randomly drawn from the population differ in type.

#### Definition 1.

The *expected heterozygosity* in a population for a given locus with *J* distinct alleles is defined as 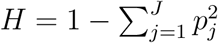, where *p*_*j*_ is the frequency of allelic type *j*.

We denote by *H*_*k*_ the expected heterozygosity of source population *k* at a locus. We have 0 ⩽ *H*_*k*_ < 1, with *H*_*k*_ = 0 if and only if source population *k* has only a single allelic type of nonzero frequency. We refer to expected heterozygosity simply as heterozygosity.

### 2.2 Fixation index

The fixation index *F*_*ST*_ is a measure of genetic divergence among a set of subpopulations. In its general form, it is computed from *H*_*S*_, the mean of the heterozygosities of the subpopulations, and *H*_*T*_, the heterozygosity of a population formed by pooling the subpopulations into a single “total” population.

#### Definition 2.

The *fixation index, F*_*ST*_ is defined as *F*_*ST*_ = (*H*_*T*_ − *H*_*S*_)*/H*_*T*_, where *H*_*T*_ is the heterozygosity of the total population and *H*_*S*_ is the mean heterozygosity across subpopulations.

The fixation index can be regarded as a measure of genetic divergence between two populations, with *F*_*kℓ*_ denoting the value of *F*_*ST*_ between source populations *k* and *ℓ*. We assume that the two subpopulations have the same contribution to the overall population, so that they are weighted equally in producing the total population. We also assume that when pooled together, they produce a polymorphic population. In other words, we disallow the case in which there is some allelic type *j* for which *p*_*kj*_ = *p*_*ℓ j*_ = 1.

For this pairwise scenario, 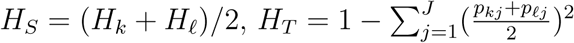, and

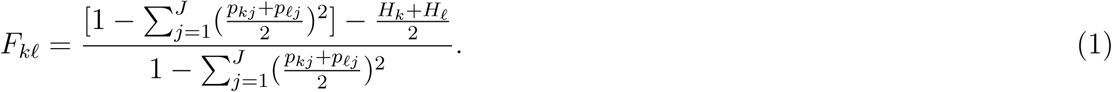

We can observe by the Cauchy-Schwarz inequality that 0 ⩽ *F*_*kℓ*_ ⩽ 1, with *F*_*kℓ*_ = 0 requiring *p*_*kj*_ = *p*_*ℓ j*_ for all *j. F*_*kℓ*_ = 1 requires *H*_*S*_ = *H*_*k*_ = *H*_*ℓ*_ = 0.

### 2.3 Allele frequency dot product

We will have occasion to use a quantity, *C*_*kℓ*_, the probability that, when randomly drawing one allele from population *k* and one allele from population *ℓ*, the two alleles differ in type. For population *k*, let *<u>p</u>*_*<u>k</u>*_ denote a *J* × 1 column vector of its allele frequencies. *C*_*kℓ*_ can then be written as 1 minus the dot product of the allele frequency vectors of populations *k* and *ℓ*:

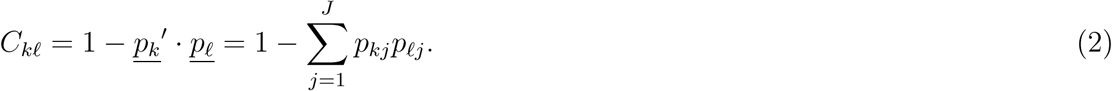

Note that this quantity can be viewed as a generalization of heterozygosity to two populations, as *H*_*k*_ = *C*_*kk*_. Because we exclude the case in which populations *k* and *ℓ* are fixed for the same allelic type, *C*_*kℓ*_ strictly exceeds 0, so that 0 < *C*_*kℓ*_ ⩽ 1. The upper bound of 1 is achieved if populations *k* and *f* share no allelic types in common.

We can rewrite eq. 1 as

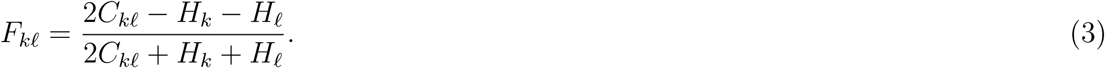

If *F*_*kℓ*_ < 1, then we can solve for *C*_*kℓ*_:

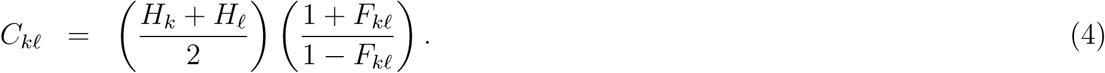

Recall that *F*_*kℓ*_ = 1 implies *H*_*k*_ = *H*_*ℓ*_ = 0, so that populations *k* and *ℓ* each have only a single allelic type with nonzero frequency. We have excluded the case in which the two populations are fixed for the same allelic type; hence, they must be fixed for different allelic types, and *C*_*kℓ*_ = 1.

By the Cauchy-Schwarz inequality, 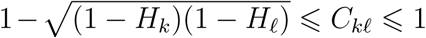 (Mehta *et al.*, 2019, eq. 7). Equality in the lower bound requires *p*_*kj*_ = *p*_*ℓ j*_ for all *j*, and hence *H*_*k*_ = *H*_*ℓ*_. Rewriting this inequality with eq. 4, we obtain the allowable space of *F*_*kℓ*_ given *H*_*k*_, *H*_*ℓ*_ ∈ [0, 1):

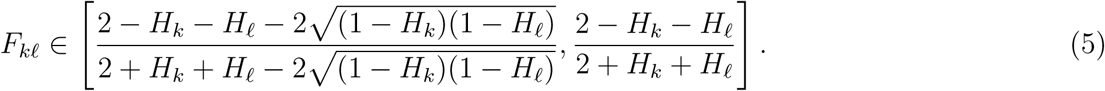

The lower limit is achieved if and only if the two populations *k* and *ℓ* are identical, with *H*_*k*_ = *H*_*ℓ*_ and *p*_*kj*_ = *p*_*ℓj*_ for all *j*. The upper limit is achieved if and only if populations *k* and *f* share no allelic types in common.

Appendix A of Mehta *et al.* (2019) shows that given *H*_*k*_ and *H*_*ℓ*_ in [0, 1), if the number of distinct alleles *J* is not fixed, then we can choose allele frequency vectors *<u>p</u>*_*<u>k</u>*_ and *<u>p</u>*_*<u>ℓ</u>*_ such that each *C*_*kℓ*_ in value in 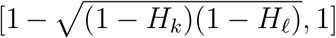 is achievable. The lower bound is achievable only if *H*_*k*_ = *H*_*ℓ*_. Hence, each value in the interval in eq. 5 for *F*_*ST*_ is also achievable by some pair of allele frequency vectors *<u>p</u>*_*<u>k</u>*_ and *<u>p</u>*_*<u>ℓ</u>*_, the lower bound only if *H*_*k*_ = *H*_*ℓ*_.

### 2.4 Admixture model

In our *K*-source-population model, *K* ⩾ 2, we follow Section 2.2 in assuming that no two populations are fixed for the same allelic type. We now make a stronger assumption that no two populations are identical, so that for each (*k, ℓ*), some *j* exists for which *p*_*kj*_ ≠ *p*_*ℓj*_.

Following a commonly used approach for describing variation in an admixed population, we treat allele frequencies in the admixed population as linear combinations of those of the source populations (e.g. Pritchard *et al.*, 2000; Boca & Rosenberg, 2011). It is convenient to assume that no source population can have its vector of allele frequencies written as the linear combination of vectors of allele frequencies of other source populations; otherwise, an admixed population would not have a unique representation as a linear combination of sources. We thus assume that not only are no two source populations identical, no source population can be described as an admixture of two or more of the other sources.

Note that the assumption that no population is a linear combination of the others also excludes linear combinations with one or more negative coefficients. In addition, because the maximal number of vectors of length *J* that can be linearly independent is *J*, the assumption implies that *J* ⩾ *K*. A succinct way of describing the linear independence assumption is that if we define the *J* × *K* matrix of allele frequencies in the source populations,

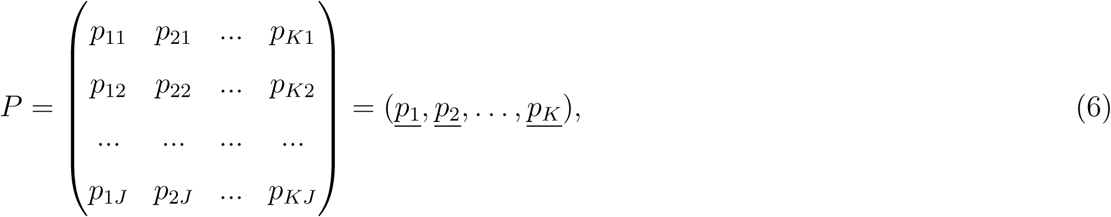

then we assume that *P* has rank *K*.

For the admixed population generated from the *K* source populations, we denote by *γ*_*k*_ the admixture fraction for source population *k*, so that for each *k* with 1 ⩽ *k* ⩽ *K*, fraction *γ*_*k*_ of the ancestry of the admixed population, 0 ⩽ *γ*_*k*_ ⩽ 1, derives from source population *k*. We denote by *<u>γ</u>* the *K* × 1 column vector of admixture fractions. This vector represents a point in the simplex Δ^*K*−1^, the set of all vectors of *K* nonnegative entries with 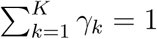.

The frequency of allele *j* in the admixed population is denoted by 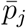. According to the linear combination assumption, we have

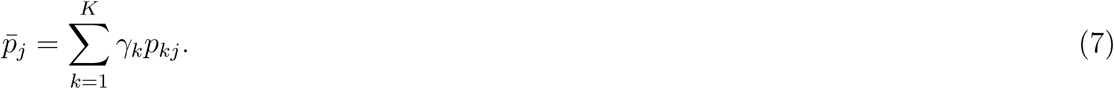

## 3 General case: *K* source populations

Our goal is to study the heterozygosity of the admixed population. Using Definition 1 with eq. 7, we compute the heterozygosity for the admixed population, which we denote by *H*_adm_:

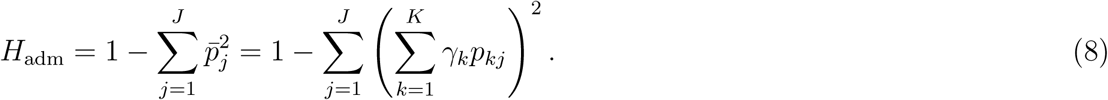

The heterozygosity of the admixed population can be written in terms of the heterozygosities of the source populations and the dot products of the allele frequencies. Using eq. 4 and the formula for *H*_adm_ in eq. 8, we have:

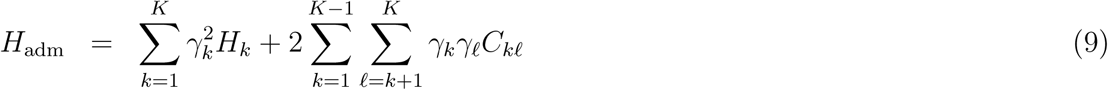

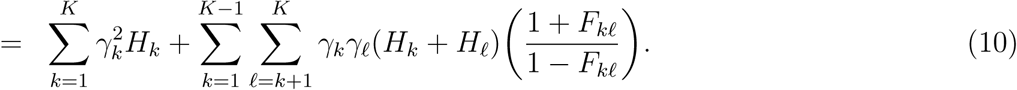

The last simplification can be made only for *F*_*kℓ*_ ≠ 1; if *F*_*kℓ*_ = 1, then eq. 9 is used, or, as noted after eq. 4, (*H*_*k*_ + *H*_*ℓ*_)(1 + *F*_*kℓ*_)/(1 − *F*_*kℓ*_) is understood to equal 2.

With the formula for *H*_adm_ established, we now explore how *H*_adm_ varies in relation to the admixture fractions <u>*γ*</u>. Given the allele frequencies *P*, we determine how small and how large *H*_adm_ can be over the space of possible values of *<u>γ</u>*. We write *H*_*m*_ for the smallest heterozygosity among the source populations, *H*_*m*_ = min_*k*∈{1,2,…,*K*}_ *H*_*k*_, and *H*_*M*_ for the largest heterozygosity among the source populations, *H*_*M*_ = max_*k*∈{1,2,…,*K*}_ *H*_*k*_.

### 3.1 Minimum of *H*_adm_ in terms of the ancestry proportions

For the minimum of *H*_adm_ over vectors (*γ*_1_, *γ*_2_,…, *γ*_*K*_), we can immediately observe from the form of eq. 10 that for a fixed set of source population allele frequencies *P, H*_adm_ is minimized as a function of the admixture fractions when the admixed population consists of only one of the source populations.

#### Proposition 3.

The minimum of *H*_adm_ as a function of the ancestry proportions *<u>γ</u>* is *H*_*m*_ = min_*k*∈{1,2,…,*K*}_ *H*_*k*_, the smallest heterozygosity among the source populations, and it is obtained when the admixed population consists solely of that source population.

*Proof.* To obtain this result, we use eq. 10 and the fact that *H*_*k*_ ⩾ *H*_*m*_ for all *k*:

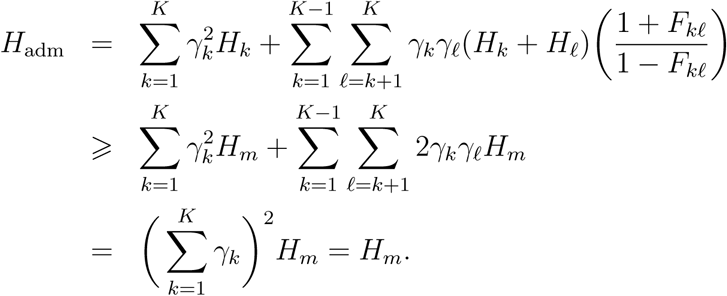

Because equality is achieved when *γ*_*m*_ = 1 and *γ*_*k*_ = 0 for all *k* ≠ *m*, we have shown that the minimal value of *H*_adm_ as a function of the ancestry proportions is *H*_*m*_. □

The result applies whether or not *H*_1_, *H*_2_,…, *H*_*K*_ are mutually distinct. If two or more of *H*_1_, *H*_2_,…, *H*_*K*_ are tied for the minimal heterozygosity *H*_*m*_, then the minimum of *H*_adm_ is achieved at each vector associated with complete ancestry from one of the minimally heterozygous populations.

A consequence of Proposition 3 is that if all *K* populations have the same heterozygosity *H*_*m*_, then *H*_adm_ > *H*_*m*_ for all ancestry vectors *<u>γ</u>* with two or more nonzero entries. In particular, note that *F*_*kℓ*_ > 0 for each (*k, ℓ*), *k* ≠ *ℓ*, by the assumption that each pair of source populations has distinct allele frequencies. Hence, (*H*_*k*_ + *H*_*ℓ*_)(1 + *F*_*kℓ*_)/(1 − *F*_*kℓ*_) > 2*H*_*m*_ for each (*k, ℓ*), *k* ≠ *ℓ*. Because at least one product *γ*_*k*_*γ*_*ℓ*_ is positive, the inequality *γ*_*k*_*γ*_*ℓ*_(*H*_*k*_ + *H*_*ℓ*_)(1 + *F*_*kℓ*_)/(1 − *F*_*kℓ*_) ⩾ 2*γ*_*k*_*γ*_*ℓ*_*H*_*m*_ is strict for at least one (*k, ℓ*), so that 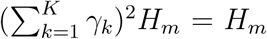. This same reasoning shows that if two or more populations are tied with heterozygosity *H*_*m*_, then *H*_adm_ > *H*_*m*_ for each *<u>γ</u>* with two or more nonzero entries.

### 3.2 Maximum of *H*_adm_ in terms of the ancestry proportions

To obtain the maximum of *H*_adm_ over the space of values of *<u>γ</u>*, we write eq. 9 as a quadratic form in terms of the ancestry proportions,

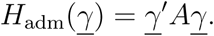

Here, *<u>γ</u>*′ represents the transpose of the column vector *<u>γ</u>* and *A* is the *K* × *K* symmetric matrix with the *H*_*k*_ on the diagonal and the *C*_*kℓ*_ off the diagonal:

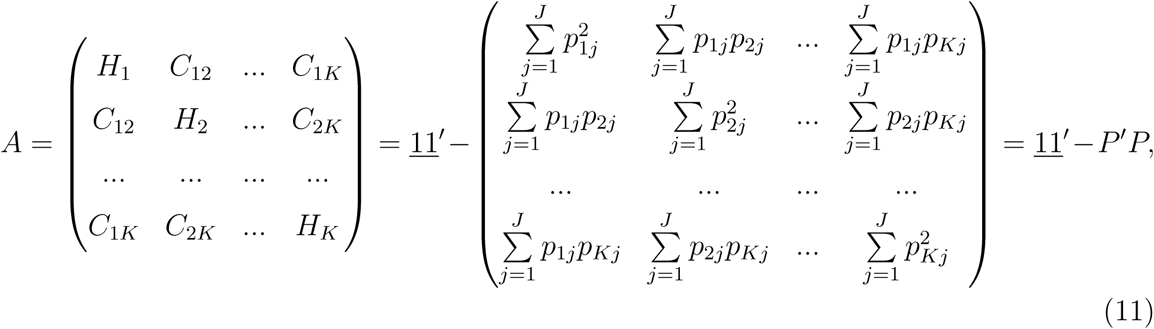

where *P* is the *J* × *K* allele frequency matrix (eq. 6) and <u>1</u> is a *K* × 1 vector of ones.

Maximizing *H*_adm_ in terms of *<u>γ</u>* is equivalent to finding 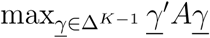 subject to <u>1</u>′*<u>γ</u>* = 1. We denote by *<u>γ</u>*<u>_arg max_</u> the location of the maximal value of *H*_adm_. We first observe that *<u>γ</u>*<u>_arg max_</u> is sometimes interior to the simplex, and that it sometimes lies at a vertex.

#### Proposition 4.

Consider the case of *K* source populations, *K* ⩾ 2.

i. There exists some collection of source population allele frequencies *P* and some collection of admixture proportions *<u>γ</u>* for which the heterozygosity of the admixed population exceeds the heterozygosity *H*_*M*_ of the most heterozygous source population.
ii. There exists some collection of source population allele frequencies *P* for which *no* collection of admixture proportions *<u>γ</u>* produces an admixed population with heterozygosity greater than the heterozygosity *H*_*M*_ of the most heterozygous source population.

*Proof.*

i. Consider *K* populations, each with different allele frequencies, but identical heterozygosity: *<u>p</u>*_*<u>k</u>*_ ≠ *<u>p</u>*_*<u>ℓ</u>*_ for *k* ≠ *ℓ* but *H*_*k*_ = *H* for *k* = 1, 2,…, *K*. Suppose that a locus has *K* + 1 distinct alleles, and that the allele frequencies are 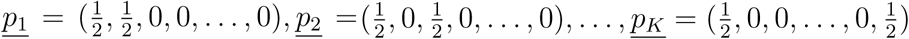. By eq. 9, 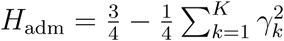, which is minimized if and only if 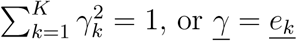 for some *k*. The minimal value of *H*_adm_ is thus 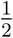, all other values of the admixture proportions resulting in 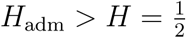.
ii. Consider *K* populations and a locus with *K* distinct alleles. Suppose that the number of distinct alleles at the locus is *k* for population *k*, with 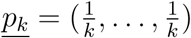. Hence, 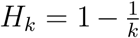 and, in particular, *H*_1_ <…< *H*_*K*_. We show that *H*_adm_ ⩽ *H*_*K*_ irrespective of *<u>γ</u>*.

By eq. 9,

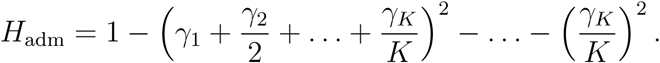

By the Cauchy-Schwarz inequality:

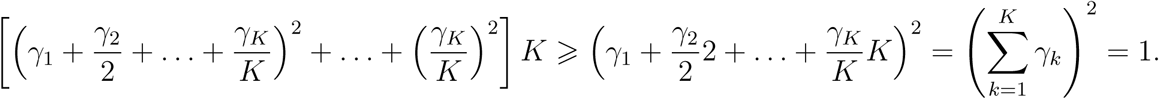

Thus, 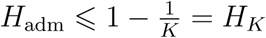. □

Note that it is trivial to see that in general, 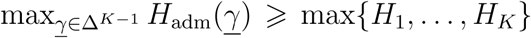: the *K* source populations simply correspond to the *K* vertices of the simplex. This result that the maximal *H*_adm_ is at least is great as the heterozygosity of the most heterozygous source population immediately implies 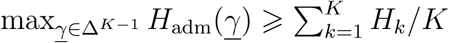.

Having established that the maximum can be at a vertex or an interior point of the simplex, we now provide a general theorem. The theorem gives the location of the maximum when it lies in the interior of Δ^*K*−1^, rather than on the boundary, assuming a condition applies on the allele frequencies. The proof is in Appendix 1.

#### Theorem 5.

Suppose that <u>1</u>′(*P*′*P*)^−1^<u>1</u> ≠ 1. Suppose also that 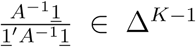. Then the maximum of *H*_adm_ as a function of the ancestry proportions *<u>γ</u>* ∈ Δ^*K*−1^ is attained at <u>*γ*_arg max_</u> = *<u>γ</u>*\*, where:

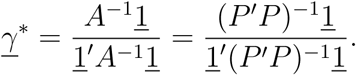

The maximum is equal to:

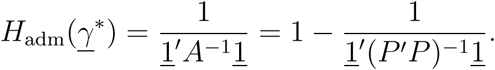

If 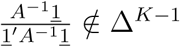, then <u>*γ*_arg max_</u> lies on the boundary of the set {*<u>γ</u>* : <u>1</u>′*<u>γ</u>* = 1 and *<u>γ</u>* ∈ Δ^*K*−1^}.

The following corollary, also proven in Appendix 1, further describes the possible locations of the maximum of *H*_adm_. Note that if the maximum is not at *<u>γ</u>*\*, then it lies at a point that has some elements equal to 0, with the nonzero subvector having a similar form to *<u>γ</u>*\*, but in a lower number of dimensions. Thus, the maximum can occur in a scenario in which the admixture involves only a strict subset of the source populations.

Consider a nonempty subset 𝒮 ⊂ {1, 2,…, *K*}. Define by *A*_𝒮_ the |𝒮| × |𝒮| matrix that has diagonal terms *H*_*k*_ for each *k* ∈ 𝒮 and off-diagonal terms *C*_*kℓ*_ for each distinct *k, f* ∈ 𝒮. Additionally, denote by *P*_𝒮_ the matrix consisting of the columns of *P* corresponding to the subset 𝒮. *P*_𝒮_ contains the allele frequencies for the source populations in 𝒮.

#### Corollary 6.

Suppose that 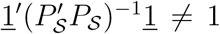 for all nonempty 𝒮 ⊂ {1, 2,…, *K*}. Then the maximum of *H*_adm_ as a function of the ancestry proportions *<u>γ</u>* ∈ Δ^*K*−1^ is attained at a point that has nonzero elements for some nonempty subset of the source populations 𝒮* ⊂ {1, 2,…, *K*}. The nonzero subvector of ancestry proportions at the location of the maximum is equal to 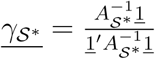.

In particular, note that *<u>γ</u>*<u>_arg max_</u> = *<u>γ</u>*\* corresponds to 𝒮* = {1, 2,…, *K*}: all source populations contribute nonzero admixture fractions. The *K* vertices of the simplex Δ^*K*−1^ correspond to the cases of 𝒮* = {*k*}, at which only one source population contributes. 𝒮 has 2^*K*^ − 1 nonempty subsets.

## 4 *K* = 2 source populations

With some general results established for the case of arbitrary *K*, we now focus on the simplest case, with *K* = 2 source populations contributing to the admixed population.

We continue to exclude the scenario in which the allele frequencies for the two source populations are identical, so that we assume *<u>p</u>*<u>_1_</u> ≠ *<u>p</u>*<u>_2_</u>. Noting that *γ*_2_ = 1 − *γ*_1_, we can consider *H*_adm_ in terms of a single admixture coefficient *γ*_1_, the admixture fraction of the first population, with *γ*_1_ ∈ [0, 1]. Using eqs. 9 and 10 with this substitution, we obtain:

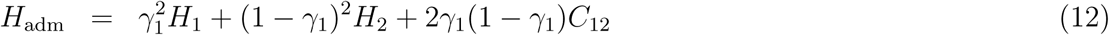

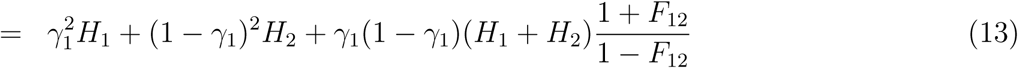

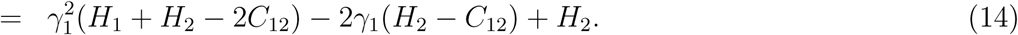

In particular, we note that from eq. 13 that *H*_adm_ is increasing as a function of *F*_12_.

From eq. 14, we can see that *H*_adm_ is concave down as a function of *γ*_1_. We have 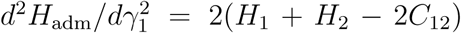. By Definition 1 and eq. 2, 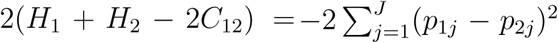. Because *<u>p</u>*<u>_1_</u> ≠ *<u>p</u>*<u>_2_</u>, *p*_1*j*_ ≠ *p*_2*j*_ for at least one choice of *j*, and hence 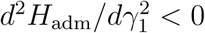. By symmetry, *H*_adm_ is also concave down as a function of *γ*_2_.

To illustrate eq. 13, for fixed values of *H*_1_ and *H*_2_, Figure 1 plots *H*_adm_ as a function of *γ*_1_ for a variety of values of *F*_12_. The figure illustrates the concave-down quadratic nature of *H*_adm_ as a function of *γ*_1_. We observe that for each value of *F*_12_ considered, the minimum of *H*_adm_ occurs at (*γ*_1_, *γ*_2_) = (0, 1), reflecting the result of Proposition 3 that the minimum occurs when the admixed population consists solely of the less heterozygous source population. In accord with the fact that in eq. 13, *H*_adm_ increases for fixed *H*_1_, *H*_2_, and *γ*_1_ with increasing *F*_12_, the value at the maximum increases with increasing *F*_12_. The location of the maximum lies at a value of 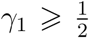, decreasing with increasing *F*_12_. This location has a pattern where for larger values of *F*_12_, it lies interior to the unit interval, and for smaller values of *F*_12_, it occurs when the admixed population consists solely of the more heterozygous source population. We now consider this pattern in more detail.

**Figure 1:**
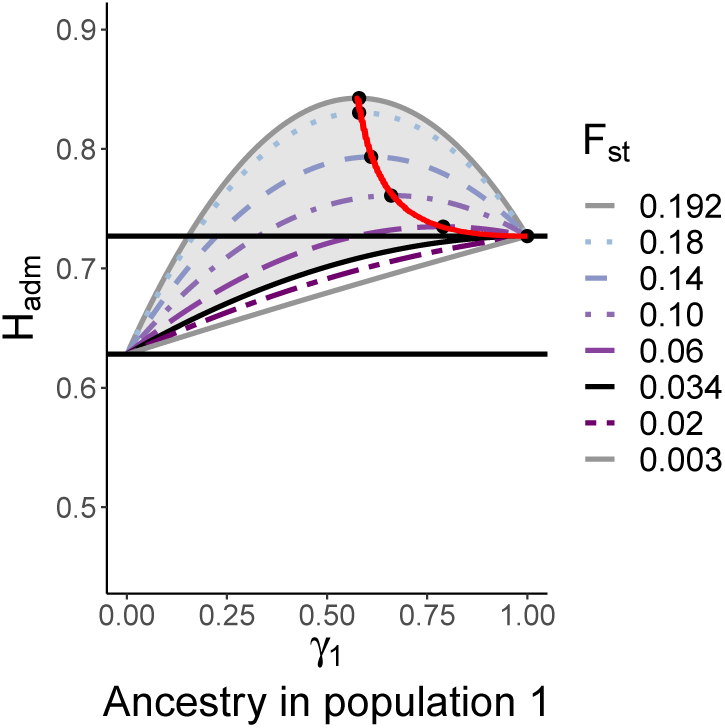
*H*_adm_ versus *γ*_1_ for fixed values of *H*_1_ and *H*_2_. We choose *H*_1_ = 0.727 and *H*_2_ = 0.628; the horizontal lines represent *H*_adm_ = *H*_1_ and *H*_adm_ = *H*_2_. Eq. 13 is plotted for multiple values of *F*_12_, considering the allowable range of *F*_12_ values in [0.003, 0.192] as specified by eq. 5. The red curve, which plots (*γ*_1_, *H*_adm_) in terms of *H*_1_, *H*_2_, and *F*_12_ in the form of eqs. 24 and 25, indicates the maxima of *H*_adm_ as *F*_12_ varies, with black dots specifying the maxima for the specific plotted values of *F*_12_. The shaded region corresponds to the region where 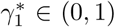, as specified by Corollary 8; the value *F*_12_ ≈ 0.034 gives the boundary of this region. The values chosen for *H*_1_ and *H*_2_ are, respectively, the mean heterozygosities across 8 European and 29 Native American populations, based on population-wise estimates in Table S20 of Pemberton *et al.* (2013). The value of *γ*_1_ can be viewed as the fraction of European ancestry in an admixed population and 1 − *γ*_1_ can be considered the fraction of Native American ancestry.

### 4.1 Minimum and maximum of *H*_adm_ in terms of the ancestry proportions

Applying the general results from Section 3.1 describing the minimum and maximum of *H*_adm_ as a function of *<u>γ</u>*, by Proposition 3, the minimum of *H*_adm_ is simply min{*H*_1_, *H*_2_}. As shown in the following proposition, the maximum can occur in one of three locations.

#### Proposition 7.

Consider two source populations with distinct allele frequencies, *<u>p</u>*<u>_1_</u> ≠ *<u>p</u>*<u>_2_</u>. As a function of *γ*_1_, *H*_adm_ is maximized at 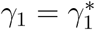, where 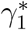 takes one of three forms.

i. If *H*_1_ < *C*_12_ and *H*_2_ < *C*_12_, then 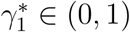 satisfies

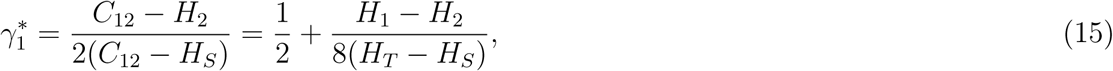

and *H*_adm_ has maximum equal to

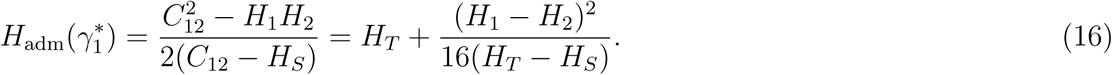
ii. If *H*_1_ < *C*_12_ and *H*_2_ ⩾ *C*_12_, then 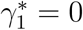 and *H*_adm_ has maximum *H*_2_.
iii. If *H*_1_ ⩾ *C*_12_ and *H*_2_ < *C*_12_, then 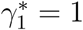 and *H*_adm_ has maximum *H*_1_.

An elementary proof appears in Appendix 2. We can see that these three cases capture all possible values of (*H*_1_, *H*_2_, *C*_12_). By the Cauchy-Schwarz inequality, (1−*C*_12_)^2^ ⩽ (1−*H*_1_)(1− *H*_2_), with equality requiring *<u>p</u>*<u>_1_</u> = *<u>p</u>*<u>_2_</u>. Hence, with *<u>p</u>*<u>_1_</u> ≠ *<u>p</u>*<u>_2_</u> assumed, either 1 − *C*_12_ < 1 − *H*_1_ and 1 − *C*_12_ ⩾ 1 − *H*_2_ (case ii), 1 − *C*_12_ < 1 − *H*_2_ and 1 − *C*_12_ ⩾ 1 − *H*_1_ (case iii), or both 1 − *C*_12_ < 1 − *H*_1_ and 1 − *C*_12_ < 1 − *H*_2_ (case i).

Note that the locations specified in Proposition 7 accord with those in Theorem 5 and Corollary 6. For *K* = 2,

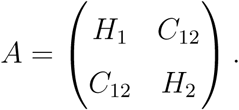

The result of Theorem 5 gives 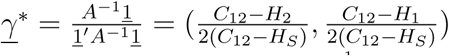. The locations in Corollary 6 are 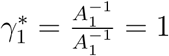 and 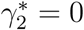, and 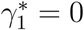 and 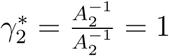.

In accord with the observation in Figure 1 that the maximal *H*_adm_ lies at a value of 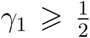 in an example with *H*_1_ ⩾ *H*_2_, the proposition demonstrates 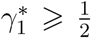 if and only if *H*_1_ ⩾ *H*_2_. To obtain this result, suppose *H*_1_ ⩾ *H*_2_. If case (i) applies, then because *H*_*T*_ > *H*_*S*_, 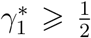. Case (ii) cannot apply because *H*_1_ < *C*_12_, *H*_2_ ⩾ *C*_12_, and *H*_1_ *H*_2_ cannot hold simultaneously. In case (iii), 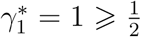. For the reverse direction, if *H*_1_ < *H*_2_ and case (i) or case (ii) applies, then 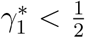. Case (iii) cannot apply because *H*_1_ ⩾ *C*_12_, *H*_2_ < *C*_12_, and *H*_1_ < *H*_2_ cannot hold simultaneously.

We also have 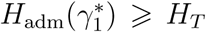 in all three cases, and 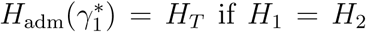. To obtain this result, we see that 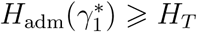 in case (i). In case (ii), 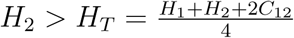 because *H*_2_ > *H*_1_ and *H*_2_ ⩾ *C*_12_. In case (iii), *H*_1_ > *H*_*T*_ because *H*_1_ > *H*_2_ and *H*_1_ ⩾ *C*_12_. Note that if *H*_1_ = *H*_2_, then case (i) applies, producing 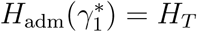.

We can succinctly describe the region where 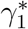 lies interior to (0, 1).

#### Corollary 8.

Consider two source populations with distinct allele frequencies, 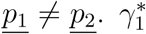 lies in (0, 1) if and only if the following inequality holds:

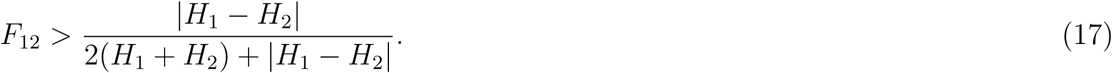

This corollary is proven in Appendix 2. Note that if *H*_1_ + *H*_2_ is fixed, then the right-hand side of eq. 17 increases with |*H*_1_ − *H*_2_|, from a minimum of 0 when *H*_1_ = *H*_2_ to a supremum of 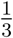 as |*H*_1_ − *H*_2_| approaches *H*_1_ + *H*_2_. Thus, in accord with the observation in Section 3.1 that *H*_adm_ > *H* for all nontrivial admixtures of equal-heterozygosity source populations, the maximal *H*_adm_ exceeds max{*H*_1_, *H*_2_} over a broader range of *F*_12_ values if |*H*_1_ − *H*_2_| is small rather than large. Moreover, if 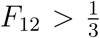, then eq. 17 necessarily holds. Hence, irrespective of *H*_1_ and *H*_2_, if the source populations are distant enough that 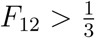, then the maximal heterozygosity exceeds the heterozygosities of the source populations.

### 4.2 Special case of *J* = 2 alleles

In the case with *K* = 2 source populations in which the locus has only *J* = 2 allelic types, it is possible to make further simplifications, as the results can be stated in terms of frequencies of one specific allele. We substitute *p*_12_ = 1 − *p*_11_ and *p*_22_ = 1 − *p*_21_.

#### Proposition 9.

Consider two source populations with distinct allele frequencies, *<u>p</u>*<u>_1_</u> ≠ *<u>p</u>*<u>_2_</u>. For a biallelic locus, *H*_adm_ is maximized at 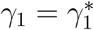, where 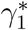 takes one of three forms.

i. If 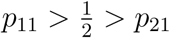 or 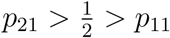, then 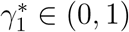 satisfies

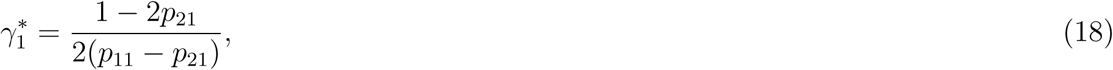

and *H*_adm_ has maximum equal to

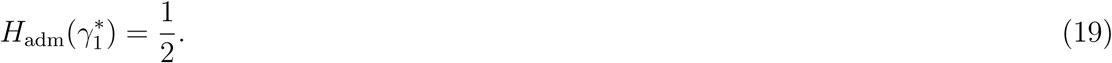
ii. If 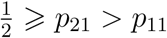 or 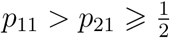, then 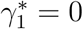 and *H*_adm_ has maximum *H*_2_.
iii. If 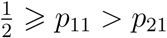 or 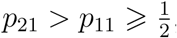, then 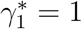 and *H*_adm_ has maximum *H*_1_.

*Proof.* We apply Proposition 7 with *J* = 2. Substituting *p*_12_ = 1 − *p*_11_ and *p*_22_ = 1 − *p*_21_ in eqs. 15 and 16, we obtain *C*_12_ − *H*_2_ = (*p*_11_ − *p*_21_)(1 − 2*p*_21_), *C*_12_ − *H*_1_ = (*p*_21_ − *p*_11_)(1 − 2*p*_11_), *C*_12_ − *H*_*S*_ = (*p*_11_ − *p*_21_)^2^, and 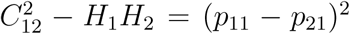, Thus, because *p*_11_ = *p*_21_ is not permitted, the quantities in eqs. 15 and 16 reduce to those of eqs. 18 and 19, respectively.

To complete the application of Proposition 7 to *K* = 2, note that case (i) of Proposition 7 occurs when (*p*_11_ − *p*_21_)(1 − 2*p*_21_) > 0 and (*p*_21_ − *p*_11_)(1 − 2*p*_11_) > 0. The first of this pair of inequalities requires both *p*_11_ − *p*_21_ > 0 and 1 − 2*p*_21_ > 0, so that *p*_11_ > *p*_21_ and 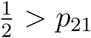, or both *p*_11_ − *p*_21_ < 0 and 1 − 2*p*_21_ < 0, so that *p*_11_ < *p*_21_ and 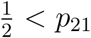. The second inequality requires both *p*_21_ − *p*_11_ > 0 and 1 − 2*p*_11_ > 0, so that *p*_21_ > *p*_11_ and 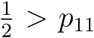, or both *p*_21_ − *p*_11_ < 0 and 1 − 2*p*_11_ < 0, so that *p*_21_ < *p*_11_ and 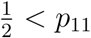. Thus, the conditions of case (i) of Proposition 7 obtain if and only if 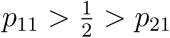 or 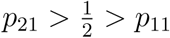.

Similarly, using the expressions for *H*_1_, *H*_2_, and *C*_12_ when *K* = 2, the conditions of case

(ii) of Proposition 7 are equivalent to 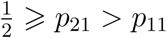 or 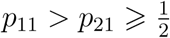. The conditions of case

(iii) are equivalent to 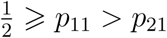 or 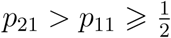.

The unit square representing the possible values of the location of the maximum appears in Figure 2. The square has six nonoverlapping regions: in Proposition 9, each of the three cases corresponds to two disjoint subsets of [0, 1]^2^. A smooth gradient exists for the regions in case (i). However, an abrupt transition occurs at the line *p*_21_ = *p*_11_ between the case-(ii) regions where 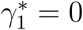 and the case-(iii) regions where 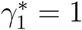. Note that the *p*_21_ = *p*_11_ line is disallowed, as it corresponds to the two populations having the same allele frequencies.

**Figure 2:**
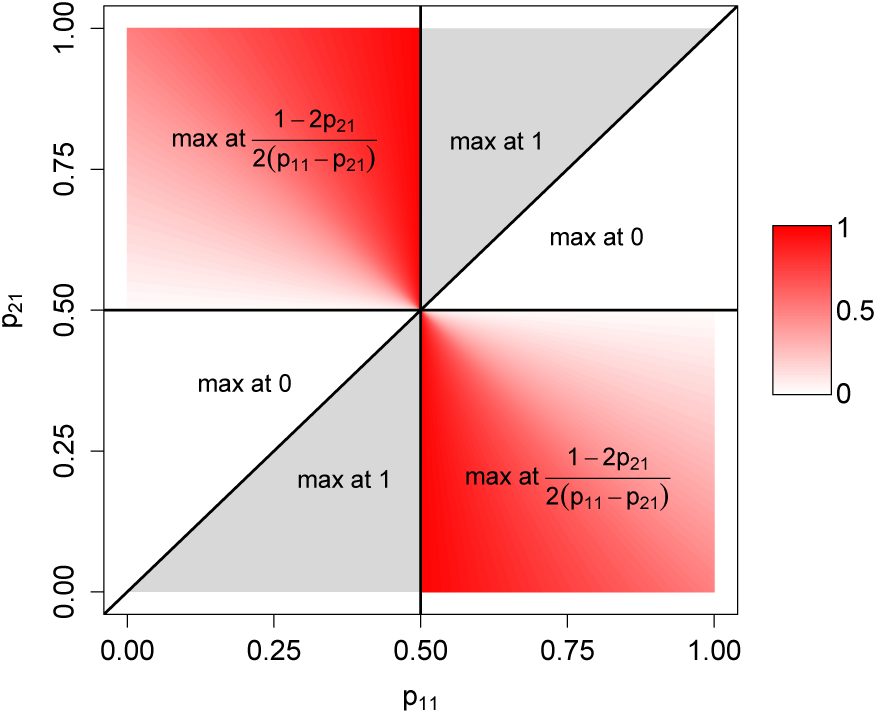
The admixture coefficient *γ*_1_ that maximizes *H*_adm_ in the case of *K* = 2 source populations and *J* = 2 allelic types. The plot shows the unit square for (*p*_11_, *p*_21_). In the red regions, the maximizing value of *γ*_1_ lies in (0, 1), whereas in the white and gray regions, it lies on one or the other boundary. The figure depicts the result of Proposition 9.

## 5 Simulations

We illustrate a number of properties of *H*_adm_ by simulating population sets for different values of *K* and *J*. Given a value of *K*, we generated allele frequency vectors for the *K* source populations from independent and identically distributed symmetric multivariate *J* -dimensional Dirichlet distributions with a common concentration parameter *α* = 1. This distribution corresponds to a uniform distribution on the simplex Δ^*J*−1^. Mathematical results concerning *H*_adm_ under the Dirichlet distribution on allele frequencies appear in Appendix 3.

First, for *K* = 2 and *K* = 3, we assessed the probability that the maximal *H*_adm_ over possible admixture vectors *<u>γ</u>* occurs interior to the simplex Δ^*K*−1^, rather than on its boundary. This computation gives the probability that the heterozygosity-maximizing admixture vector contains nonzero contributions from all *K* source populations. We considered 2 ⩽ *J* ⩽ 30 for *K* = 2 and 3 ⩽ *J* ⩽ 30 for *K* = 3, recalling the condition *J* ⩾ *K* for the *K* allele frequency vectors to be linearly independent.

For each (*K, J*), we ran 10,000 simulation replicates. In each replicate, to determine the location of the maximum, we applied Theorem 5 and Corollary 6 to identify the locations specified for each choice 𝒮 of the nonempty subset of the *K* populations with nonzero allele frequencies. Among these 2^*K*−1^ locations, excluding those outside the simplex Δ^*K*−1^, we identified the point with the largest *H*_adm_. Note that in each replicate, we observed that the 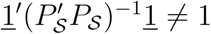 condition of Corollary 6 was satisfied for each 𝒮.

Figure 3 finds that, for both *K* = 2 and *K* = 3, the maximum of *H*_adm_ is increasingly likely to be in the interior of the simplex as the number of distinct alleles, *J*, increases. For *K* = 3, we also observe that the probability that *H*_adm_ is maximized on an edge, corresponding to nonzero contributions from two of the three source populations, exceeds the probability that it is maximized at a vertex, with only one contributing source population.

**Figure 3:**
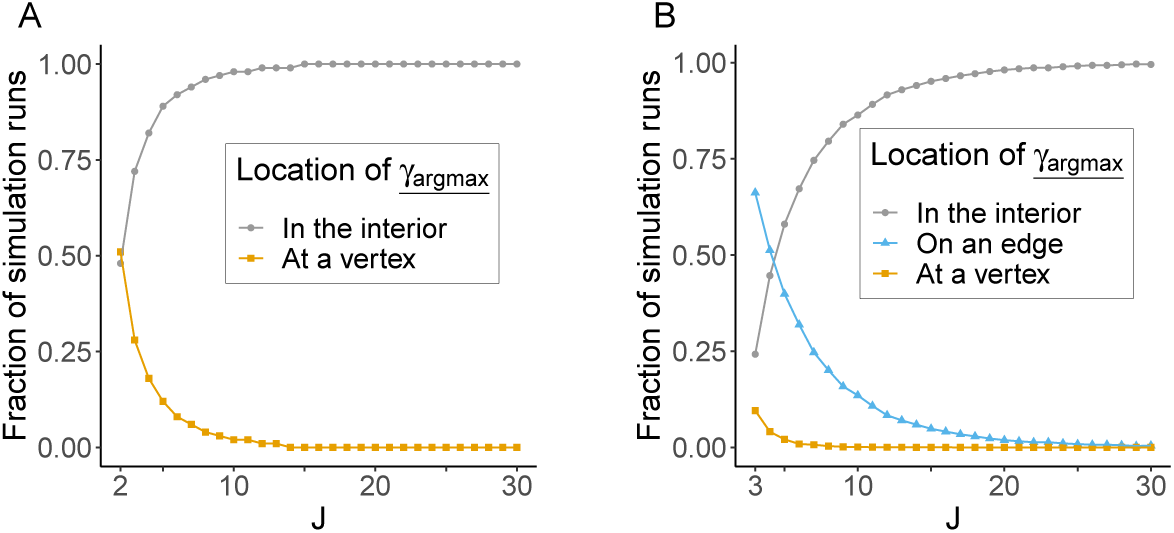
Location of the maximum of *H*_adm_ in simulation replicates. (A) *K* = 2. (B) *K* = 3. The location *<u>γ</u>*<u>_arg max_</u> can be in the interior of the simplex Δ^*K*−1^, corresponding to nontrivial admixture of all source groups, or on the boundary of the simplex. For *K* = 3, it can be on an edge, corresponding to admixture of two of three source populations, and for both *K* = 2 and *K* = 3, it can be at a vertex, corresponding to membership in only one source population. For each (*K, J*), points plotted are based on 10,000 simulations with independently and identically distributed Dirichlet-(1, 1, …, 1) distributions for the allele frequency vectors *<u>p</u>*_*<u>k</u>*_ in the *K* populations.

Next, we assessed the probability ℙ[*H*_adm_ > max{*H*_1_,…, *H*_*K*_}] in a scenario in which boththe allele frequency vectors *<u>p</u>*_*<u>k</u>*_ and the admixture fractions *<u>γ</u>* were chosen from independent Dirichlet distributions. We simulated the *<u>p</u>*_*<u>k</u>*_ as before, additionally simulating *<u>γ</u>* from a *K*- dimensional symmetric Dirichlet-(1, 1,…, 1) distribution. For each (*K, J*) with *K* = 2, 3, 4, 5 and *J* = 2, 3,…, 30, we simulated 50,000 replicate populations. Note that here, unlike in Section 2.4, we impose no restrictions on linear combinations of allele frequency vectors from the source populations, so that it is not necessarily true that *J* ⩾ *K*.

The fraction of replicates with ℙ[*H*_adm_ > max{*H*_1_,…, *H*_*K*_}] appears in Figure 4. We see that this fraction increases as *K* increases, indicating that for an admixture involving more populations, the probability is larger that the admixed population has greater heterozygosity than all source populations. This probability also increases with increasing *J*.

**Figure 4:**
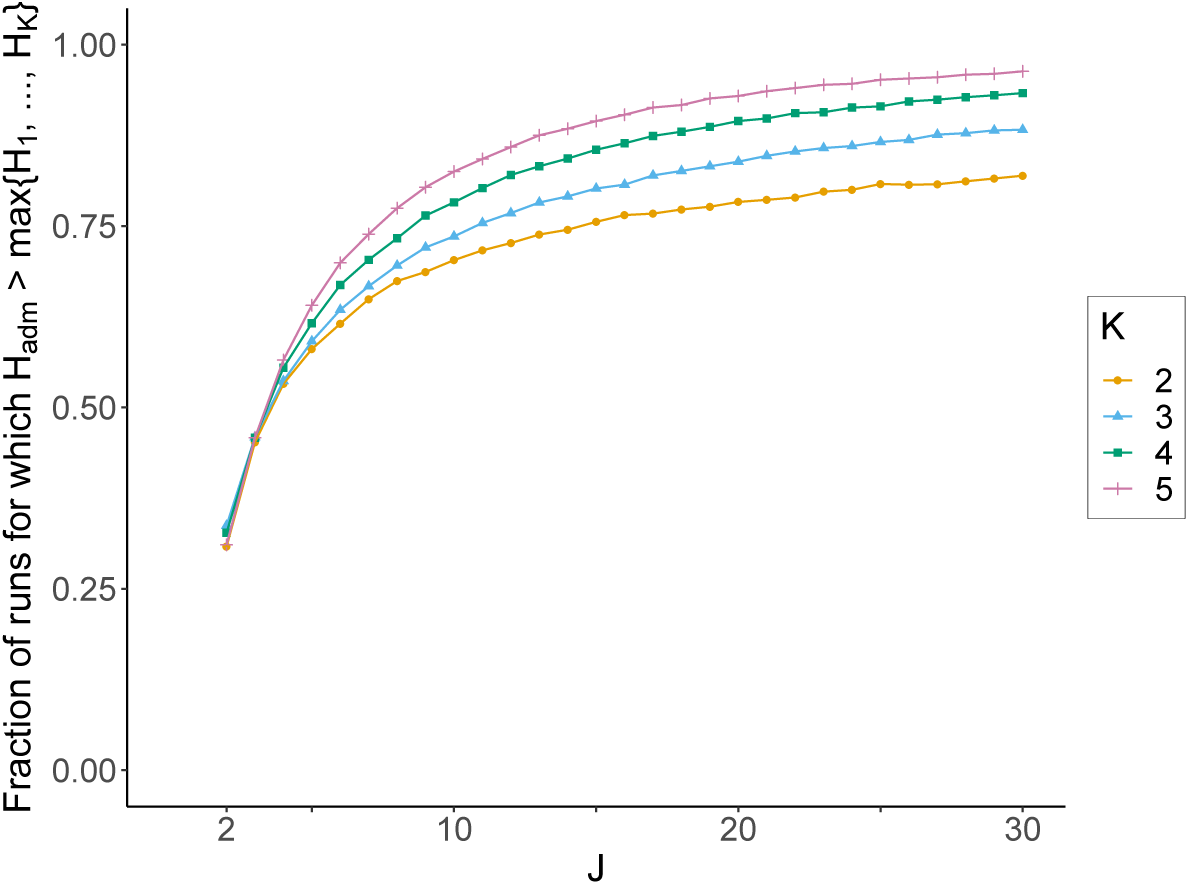
The fraction of simulation replicates for which *H*_adm_ > max {*H*_1_, …, *H*_*K*_}, for various values of *K* and *J*. For each (*K, J*), points plotted are based on 50,000 simulation replicates with independently and identically distributed Dirichlet-(1, 1, …, 1) distribitions for the allele frequency vectors *<u>p</u>*_*<u>k</u>*_ in the *K* populations, and a Dirichlet-(1, 1, …, 1) distribution for the admixture coefficient vector *<u>γ</u>*.

For the special case of *K* = 2 and *J* = 2, Proposition 15 in Appendix 3 obtains the probability analytically, ℙ[*H*_adm_ > max{*H*_1_, *H*_2_}] = 1 − log 2 ≈ 0.307. In accord with this result, the *K* = 2 curve in Figure 4 begins near (2, 0.307).

Figure 5 provides further detail on the behavior of *H*_adm_ in the *K* = 2 case by graphing *H*_adm_ versus *γ*_1_ for 10 simulation replicates chosen at random for each of three values of *J*. The figure illustrates that *H*_adm_ is a concave-down quadratic polynomial in *γ*_1_, as in eq. 14. Averaging across replicates, by examining the panels of the figure from left to right, we can also observe that 𝔼[*H*_adm_] increases as a function of *J*, as in Corollary 14 of Appendix 3. For *J* = 2, as in Proposition 9, the possible values of *H*_adm_ at the maximum are *H*_1_, *H*_2_, and 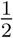.

**Figure 5:**
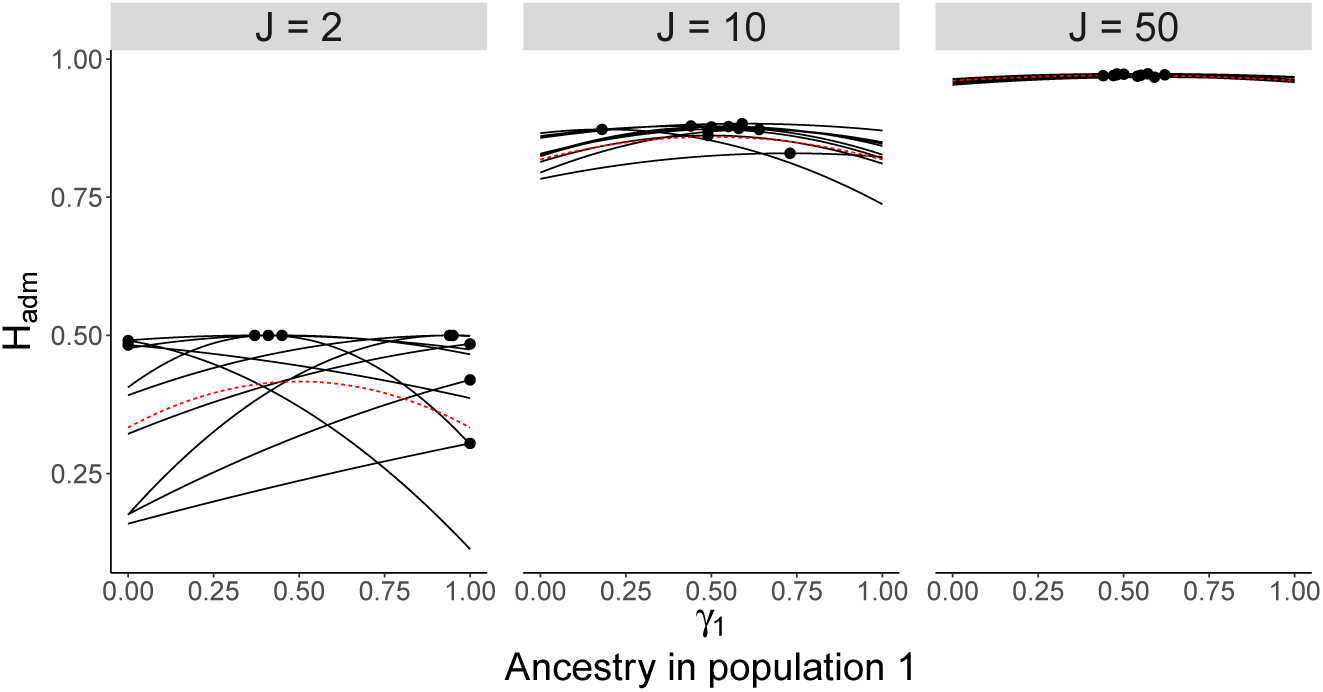
*H*_adm_ versus *γ*_1_ for 10 simulation replicates for *K* = 2 source populations, for each of three values of the number of allelic types *J*. For each replicate, allele frequency vectors *<u>p</u>*_*<u>k</u>*_ in the two populations are simulated according to Dirichlet-(1, 1, …, 1) distributions, and *H*_adm_ is plotted as a function of *γ*_1_ according to eq. 8. The maximum of *H*_adm_ is indicated by a black circle in each replicate. The red dashed lines represent the expected values of *H*_adm_ according to Corollary 14 in Appendix 3.

## 6 Application to data

We illustrate the mathematical results using data from human populations, following Boca & Rosenberg (2011) in considering data from Wang *et al.* (2008) on 678 microsatellite loci typed in 160 Europeans, 463 Native Americans, 123 Africans, and 249 individuals from admixed Mestizo populations. To represent admixed Mestizo populations under our model, we used sample allele frequencies for the Europeans and Native Americans as source populations in the *K* = 2 case, also including sample allele frequencies for the Africans for *K* = 3. As in Boca & Rosenberg (2011), we treated allele frequencies, heterozygosities, and *F*_*ST*_ values computed from the data as parametric values rather than estimates.

### 6.1 *K* = 2 source populations

We selected 20 loci at random from Wang *et al.* (2008) for illustration, choosing the same loci as in our study on *F*_*ST*_ and admixture (Boca & Rosenberg, 2011). Treating *γ*_1_ as the fraction of European ancestry and 1 − *γ*_1_ as the fraction of Native American ancestry in an admixed population, for each locus, the plot for *H*_adm_ versus *γ*_1_ appears in Figure 6. Following Proposition 3, the minimal value of *H*_adm_ lies either at *γ*_1_ = 0 or at *γ*_1_ = 1 for all the loci. For 12 of the 20 loci, the maximum of *H*_adm_ lies in the interior of the unit interval for *γ*_1_. Seven loci have the maximum at *γ*_1_ = 1, representing membership in the more heterozygous European population, and only one locus has the maximum at *γ*_1_ = 0, representing membership in the less heterozygous Native American population.

**Figure 6:**
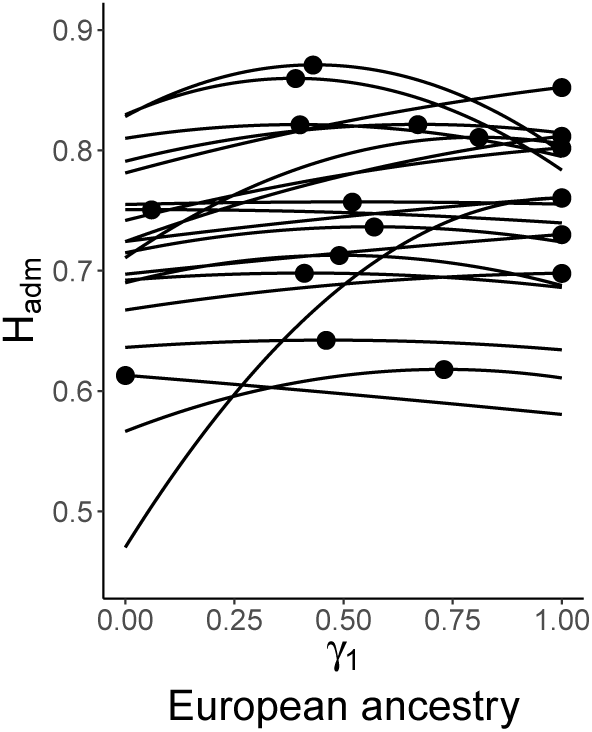
*H*_adm_ versus *γ*_1_ for 20 random loci from Wang *et al.* (2008). The two source populations providing the allele frequencies are the European and Native American populations, with *γ*_1_ corresponding to membership in the European population. *H*_adm_ is plotted according to eq. 8. Circles indicate the location of the maximum along each curve.

Examining all 678 loci, 52% have the maximum in the interior, 39% at *γ*_1_ = 1, and 8% at *γ*_1_ = 0. That more loci have the maximum at *γ*_1_ = 1 than at *γ*_1_ = 0 is expected from the fact that European populations generally tend to be have greater heterozygosity than Native American populations (e.g. Pemberton *et al.*, 2013).

The Dirichlet model in Corollary 14 in Appendix 3 and Figures 3 and 5 predicts a dependence of the location of the maximum on the number of distinct alleles of a locus, with the probability that the maximum lies in the interior increasing with the number of distinct alleles. The data produce a trend in the same direction as this prediction. The mean numbers of distinct alleles are 9.44, 10.41, and 10.74, for the loci with 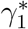 at 0, 1, and in (0, 1), respectively (one-way ANOVA, *P* = 0.01, *F* test, 2 df). The mean number of distinct alleles for the loci with the maximum on either boundary is 10.24, smaller than the mean of 10.74 for those with the mean in the interior (*P* = 0.04, two-tailed *t* test).

### 6.2 Comparison of predicted *H*_adm_ to observed *H*_adm_

Next, we compare predicted and observed *H*_adm_ values for the 678 loci for the admixed Mestizo population. In this approach, we used estimated locus-wise values of *γ*_1_ in the Mestizo population together with locus-wise heterozygosities in the European and Native American populations to “predict” locus-wise Mestizo heterozygosities. The prediction is compared to the observed heterozygosity value to examine if our formulas for the heterozygosity of an admixed population are reflected in actual heterozygosities in an admixed group.

This computation follows a similar computation of Boca & Rosenberg (2011). The estimated admixture fractions, computed for the same data, are taken from Schroeder *et al.* (2009), who obtained them by a maximum likelihood approach (Millar, 1987) that does not take into account the source population heterozygosities. Using these estimates, locus-wise heterozygosity estimates in the source populations, and locus-wise *F*_*ST*_ values calculated from the allele frequencies in the source populations, we predicted *H*_adm_ with eq. 13.

The predicted and observed *H*_adm_ values for individual loci are compared in Figure 7. In general, the observation closely matches the prediction (Figure 7A), with the correlation between the observed and predicted *H*_adm_ values equaling 0.978 (Figure 7B). For 56% of the 678 loci, the prediction provides an underestimate of the observed value.

**Figure 7:**
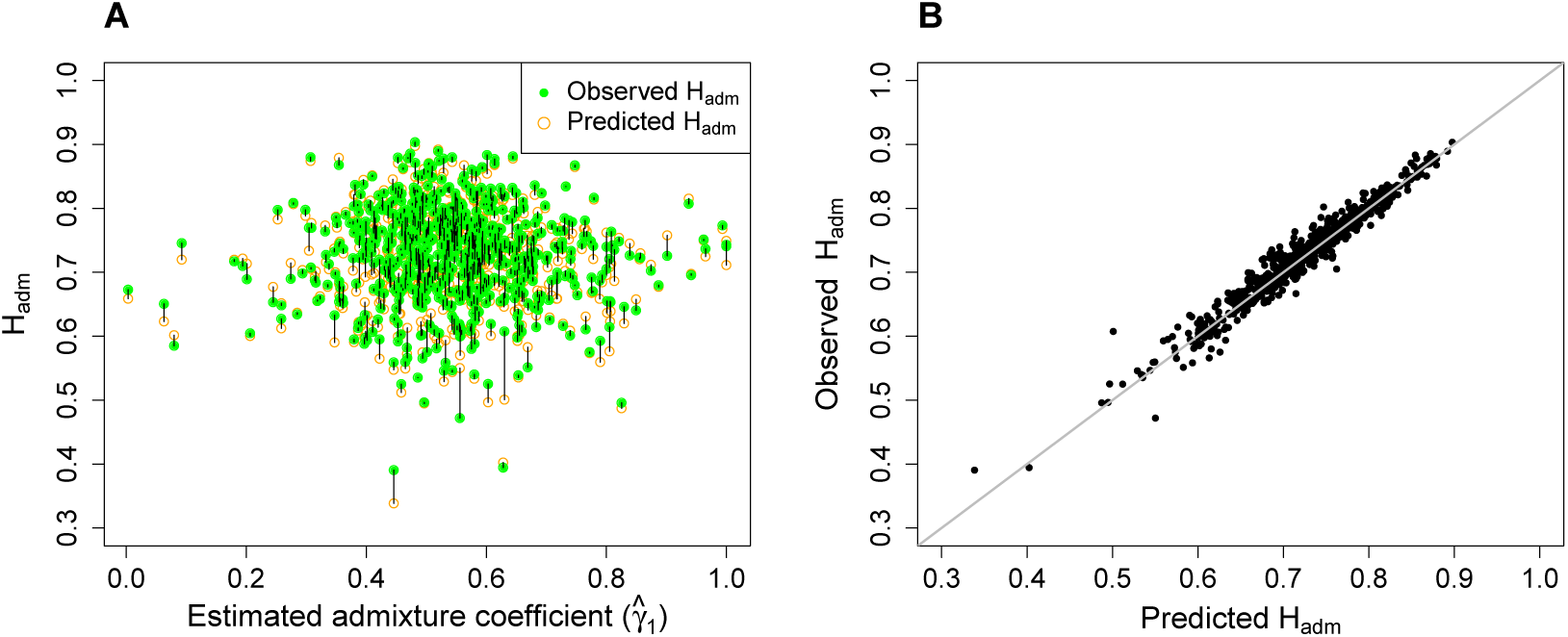
Predicted and observed *H*_adm_. (A) The predicted and observed *H*_adm_ values for an admixed Mestizo population are plotted against the locus-wise estimated European admixture fraction 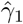 in the Mestizo population, estimated by maximum likelihood. The prediction is based on eq. 8, using European and Native American allele frequencies estimated from Wang *et al.* (2008) as *<u>p</u>*<u>_1_</u> and *<u>p</u>*<u>_2_</u>, respectively, together with the maximum likelihood estimate of *γ*_1_. The observation is based on *H*_adm_ computed from Definition 1, inserting estimated allele frequencies from Wang *et al.* (2008) for the Mestizo population. (B) The observed *H*_adm_ value is plotted against the predicted *H*_adm_ value. The identity line is shown in gray. In both panels, each point represents one of the 678 loci used. The correlation coefficient between the predicted and observed *H*_adm_ values is 0.978.

### 6.3 *K* = 3 source populations

We now consider the European, Native American, and African populations as the source populations, using *γ*_1_ for the proportion of European ancestry, *γ*_2_ for Native American ancestry, and *γ*_3_ for African ancestry. We select 3 loci for illustration, choosing the same ones as in a similar analysis of Boca & Rosenberg (2011).

Plots for *H*_adm_ over the unit simplex for (*γ*_1_, *γ*_2_, *γ*_3_) appear in Figure 8. Each plot depicts *H*_adm_ as a function of (*γ*_1_, *γ*_2_, *γ*_3_) for a specific locus. The three panels show the possible locations of the maximal value of *H*_adm_: in the first panel, the maximum lies in the interior of the simplex; in the second panel, at a vertex, and in the third panel, on an edge.

**Figure 8:**
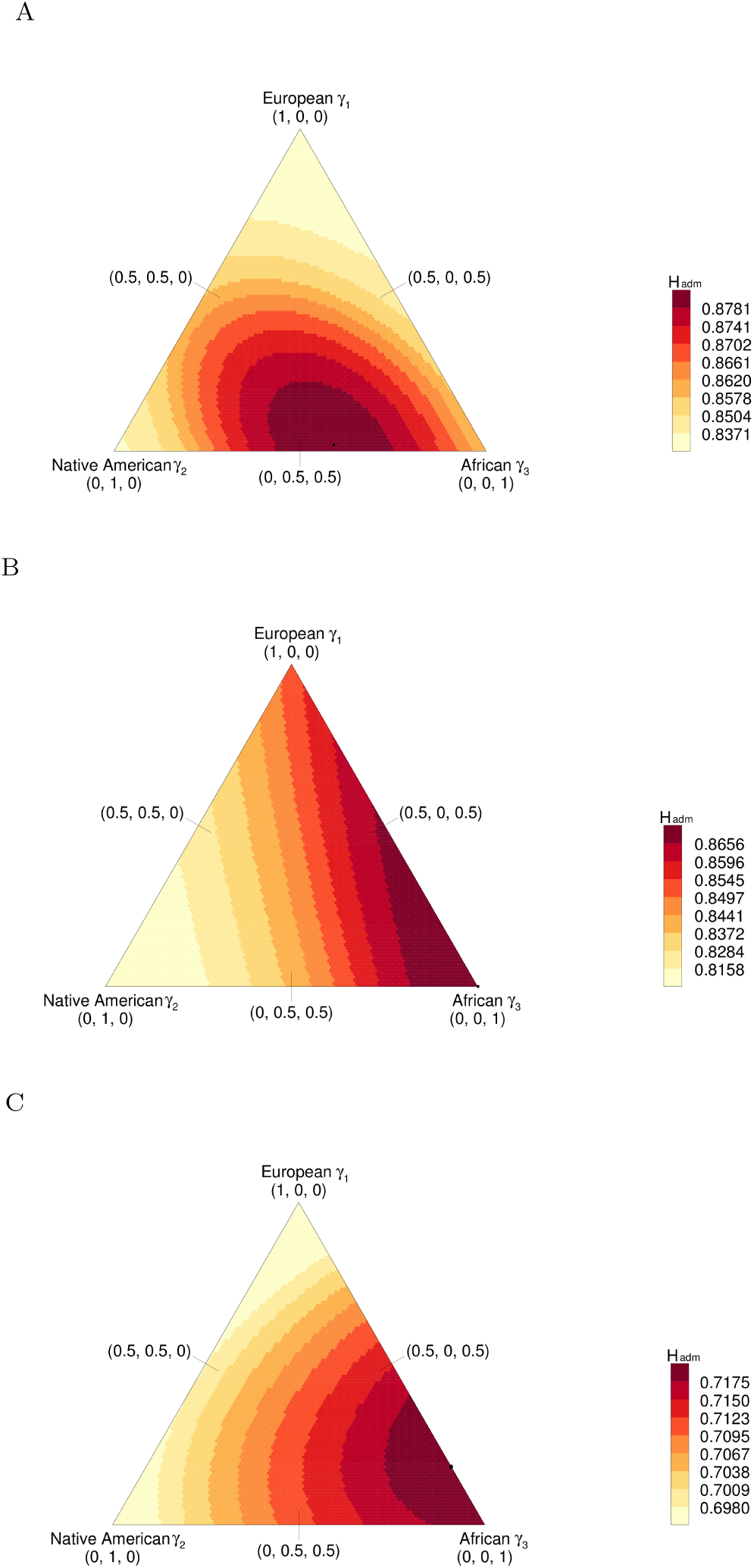
*H*_adm_ versus (*γ*_1_, *γ*_2_, *γ*_3_) for three loci. The loci are from Wang *et al.* (2008) and have 14, 14, and 8 distinct alleles, respectively. The value of *H*_adm_ is computed from eq. 8. Black circles indicate the maximum *H*_adm_. (A) Locus D2S1399: the maximum lies in the interior of the region. (B) Locus GATA101G01: the maximum lies at the (0, 0, 1) vertex. (C) Locus GATA146D07: the maximum lies on the *γ*_2_ = 0 edge.

Considering all 678 loci, 14% have the maximum in the interior of the region, with *γ*_1_ > 0, *γ*_2_ > 0, and *γ*_3_ > 0. The fractions with the maximum on an edge are 19% for a maximum on the edge with *γ*_1_ = 0, 26% on the *γ*_2_ = 0 edge, and 5% on the *γ*_3_ = 0 edge. The fractions with the maximum at a vertex are 27% for the vertex (0, 0, 1), 2% for (0, 1, 0), and 5% for (1, 0, 0). The observations that (0, 0, 1) is the vertex with the largest number of maxima and (1, 0, 1) is the edge with the most maxima accord with the fact that African populations have generally higher heterozygosity than European populations, which in turn have higher heterozygosity than Native American populations (e.g. Pemberton *et al.*, 2013).

## 7 Discussion

We have considered the heterozygosity *H*_adm_ of an admixed population in terms of the admixture fractions of the source populations, and their heterozygosities and *F*_*ST*_ values at a locus. We have derived formulas describing *H*_adm_ in relation to these quantities (eqs. 8-10). In particular, we showed that *H*_adm_ is minimized over the set of possible admixture coefficient vectors when the admixed population consists of only one of the source populations (Proposition 3); hence, an admixed population is at least as heterozygous as the least heterozygous source population. The maximal *H*_adm_ has a more complex characterization, as it has the possibility of being either greater than or equal to the heterozygosity of the most heterozygous source population (Proposition 4).

In studying the possible locations of the maximal *H*_adm_ for a fixed set of source populations, we found that the maximum can lie either in the interior of the region describing the allowable values of the admixture fractions—in which case all source populations contribute to the admixed population—or on the boundary, where one or more source populations does not contribute to the admixed population (Propositions 4-6, Figures 1-3). Simulations under a Dirichlet model for allele frequencies suggest that the maximal value of *H*_adm_ lies with increasing frequency in the interior of the allowable region as *K* and *J* increase (Figure 4).

For *K* = 2 source populations, we obtained further results, in particular showing that *H*_adm_ is a concave-down quadratic polynomial in the admixture coefficient *γ*_1_ (eqs. 12-14). We obtained an analytical expression for the maximal heterozygosity of an admixture of a specific pair of source populations in terms of *H*_1_, *H*_2_, and the *F*_*ST*_ value between the two populations (Proposition 7). For fixed values of *H*_1_, *H*_2_, and the admixture fraction *γ*_1_, *H*_adm_ is increasing as a function of *F*_*ST*_ (eq. 13, Figure 1). If *H*_1_ > *H*_2_, then the admixture fraction in source population 1 that maximizes *H*_adm_ is greater than 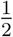 (Proposition 7), meaning that at the maximal heterozygosity of the admixed population, the contribution of the more heterozygous source population exceeds that of the less heterozygous one. Interestingly, for the *K* = 2 case with *J* = 2 allelic types, if the location of the maximal value lies in (0, 1), then the heterozygosity at the maximum is always 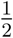 (Proposition 9 and Figure 5): irrespective of the allele frequencies of the source populations, a linear combination (*γ*_1_, *γ*_2_) always exists so that the admixed population has allele frequencies of 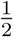 for both alleles.

For *K* = 2 source populations, a key result is that the maximal value of *H*_adm_ exceeds the larger of the two source population heterozygosities if and only if *F*_*ST*_ exceeds a bound defined by those heterozygosities (Corollary 8). Thus, with all other quantities equal, combining source populations that are more divergent rather than less divergent is more likely to lead to an admixed population with heterozygosity exceeding those of the source populations.

In human data, we observed that for heterozygosities and *F*_*ST*_ values for putative sources of Mestizo populations, the maximal *H*_adm_ was more likely to be in the interior of the unit simplex or on an edge rather than at a vertex (Figures 6 and 8). This result indicates that the heterozygosities and *F*_*ST*_ values of these populations lie in a parameter range for which admixed populations are frequently more heterozygous than all their source populations. Examining the heterozygosities of 267 worldwide populations in Table S20 of Pemberton *et al.* (2013), the 13 Mestizo populations all have heterozygosities exceeding all 29 Native American populations, and 4 have heterozygosities exceeding all 8 European populations. Interestingly, the top 10 most heterozygous populations among the 267 include all five admixed populations involving a source population from the high-heterozygosity region of Africa: a Cape Mixed Ancestry population from South Africa, and four African-American populations. Thus, our mathematical results predicting that admixed populations often exceed all their source populations in heterozygosity are reflected in admixed human groups.

For *K* = 2, our model successfully predicted the heterozygosities in an admixed population from the source population heterozygosities, the *F*_*ST*_ between the source populations, and the estimated admixture coefficient 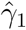 for one of the source populations (Figure 7). Because *H*_adm_ is not necessarily monotonic in the admixture fraction *γ*_1_, however, the reverse problem of using *H*_adm_ to estimate *γ*_1_ is problematic—unlike for the monotonically varying *F*_*ST*_ between an admixed population and one of the source populations (Boca & Rosenberg, 2011, Theorem 3). Given a value of *H*_adm_, source population heterozygosities *H*_1_ and *H*_2_, and *F*_*ST*_ between the source populations, two solutions to eq. 13 might exist for *γ*_1_—so that although *H*_adm_ can be predicted from *γ*_1_, it is inadvisable to proceed in the reverse direction to estimate the admixture coefficient *γ*_1_ from the heterozygosity of an admixed population.

Our approach has followed the study of *F*_*ST*_ and admixture from Boca & Rosenberg (2011), and it shares similar limitations. For example, the model assumes source population allele frequencies are known rather than estimated, and it considers only population-level rather than individual-level admixture. It also relies on patterns of variation from a single time point and does not incorporate mechanistic evolutionary processes. Despite these limitations, the observed *H*_adm_ values and those predicted under our model are strongly correlated (Figure 7B), indicating that the model captures key population features relevant to the relationship between admixture and heterozygosity. Thus, the empirical results suggest that assessing this relationship in the mathematical formulations we have presented can be useful for understanding the genetics of admixed populations.

## Acknowledgments

Support was provided by NIH grant HG005855 and NSF grant BCS-1515127.

## Appendix 1. Proofs for arbitrary *K*: Theorem 5 and Corollary 6

For the proof of Theorem 5, we first show (i) that *P*′*P* and *A* are both invertible under the conditions stated in the theorem, and that:

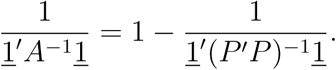

We then (ii) use constrained optimization via Lagrange multipliers to obtain the maximum of *<u>γ</u>*′*A<u>γ</u>* subject to <u>1</u>′*<u>γ</u>* = 1. This step consists of the first derivative test to find a stationary point, coupled with the second derivative test, in Lemma 10, to show that the stationary point defines a local maximum. Finally, we (iii) show that this means that the overall maximum is either at the local maximum *<u>γ</u>*\* as described in the statement of the theorem or on the boundary of the set {*<u>γ</u>* : <u>1</u>′*<u>γ</u>* = 1 aand *<u>γ</u> ∈* Δ^*K*−1^}.

*Proof of Theorem 5* (i) Because *P* is a *J* × *K* matrix with column rank *K*, the *K* × *K* matrix *P*′*P* is positive definite. As a positive definite matrix, *P*′*P* is invertible and (*P*′*P*)^−1^ is also positive definite (Graybill, 1976, pp. 21-22).

To show that *A* = <u>11</u>′ − *P*′*P* is invertible, we use the Sherman-Morrison formula for the inverse of a rank-one update of an invertible matrix (Horn & Johnson, 2012, pp. 18-19). This formula states that for an invertible square *n* × *n* matrix *X* and *n* × 1 column vectors *<u>y</u>* and *<u>z</u>, X* + *<u>yz</u>*′ is invertible if and only if 1 + *<u>z</u>*′*X*^−1^*<u>y</u>* ≠ 0, with:

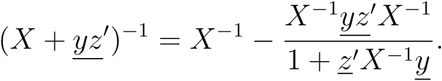

Because we assumed <u>1</u>′(*P*′*P*)^−1^<u>1</u> ≠ 1, the Sherman-Morrison formula applies with −(*P*′*P*) in the role of *X*, and *K* × 1 column vectors <u>1</u> in the role of *<u>y</u>* and *<u>z</u>. A* has inverse:

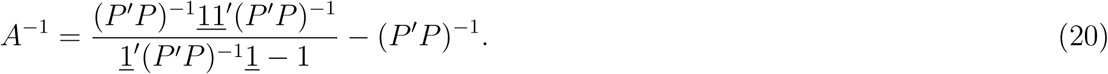

Left-multiplying by <u>1</u>′ and right-multiplying by <u>1</u>, we obtain

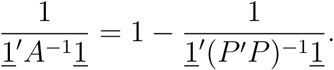

Because (*P*′*P*)^−1^ is positive definite, <u>1</u>′(*P*′*P*)^−1^<u>1</u> > 0 by definition, and because <u>1</u>′(*P*′*P*)^−1^<u>1</u> ≠1 by assumption, we conclude that 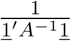 is always defined.

(ii) To maximize *<u>γ</u>* ′*A<u>γ</u>* subject to <u>1</u>′*<u>γ</u>* = 1, we use Lagrange multipliers. Let *f* (*<u>γ</u>*) = *<u>γ</u>*′ *A<u>γ</u>*, and let *g*(*<u>γ</u>*) = <u>1</u>′ *<u>γ</u>*. The Lagrange function is defined as:

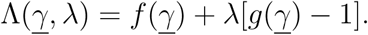

Denoting by <u>0</u> is a column vector of length *K*, we solve a system of equations for *<u>γ</u>* and *λ*,

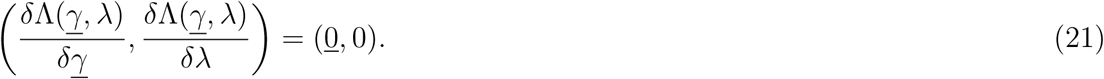

Eq. 21 includes *K* equations *δ*Λ(*<u>γ</u>, λ*)*/δγ*_*k*_ = 0 for 1 ⩽ *k* ⩽ *K*.

*A* is symmetric, so we have

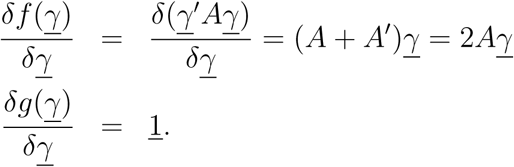

For the derivatives of the Lagrange function, we have:

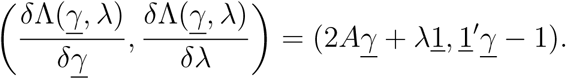

Setting the derivatives with respect to *γ* to <u>0</u> leads to:

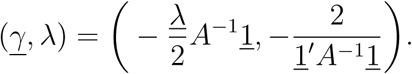

Hence, the solution for *<u>γ</u>* is:

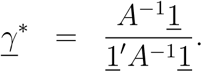

Because *<u>γ</u>*′*A<u>γ</u>* is a differentiable function of *<u>γ</u>*, its maximum on Δ^*K*−1^ can occur either on the boundary or at a critical point. The following lemma shows that the critical point 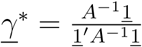 is a local maximum.

### Lemma 10.

The critical point 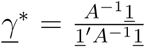 is a local maximum of *H*_adm_ seen as a function of *<u>γ</u>* on Δ^*K*−1^, under the conditions stated in Theorem 5.

*Proof.* To show that *<u>γ</u>*\* is a local maximum, we use the second derivative test for constrained optimization (e.g. Magnus & Neudecker, 2007, p. 155). This test considers the bordered Hessian matrix, representing the matrix of second derivatives of the Lagrange function Λ with respect to *λ* and the components of <u>*γ*</u>:

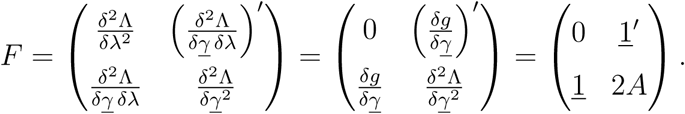

We must consider the principal minors—determinants of matrices in the upper-left corner— of *F*. We denote the upper-left corner matrix of size *r* × *r* of *F* by *F*_*r*_, for *r* = 2, 3, …, *K*. The principal minors are the det(*F*_*r*_). Using the definition of *A* from eq. 11, we obtain

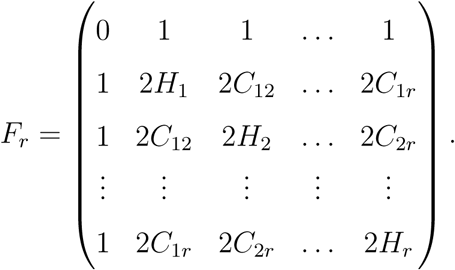

A sufficient condition for the critical point to be a local maximum is for (−1)^*r*^ det(*F*_*r*_) > 0 for each *r* (Magnus & Neudecker, 2007, p. 155). We now show that this condition is satisfied.

Using the fact that multiplying a row or column of a matrix by a scalar multiplies the determinant by that scalar, we multiply rows 2 through *r* + 1 by −1 and get

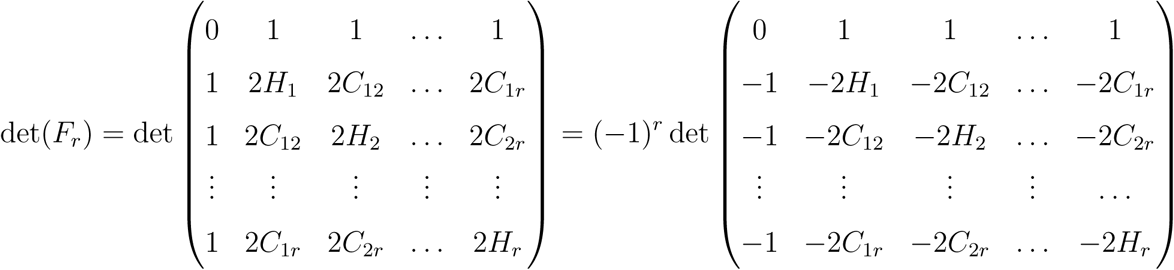

Using the fact that adding a multiple of a row or column to another row does not change the determinant, we add −2 times the first column to each of the remaining columns. We also multiply the first column by −1. We then have

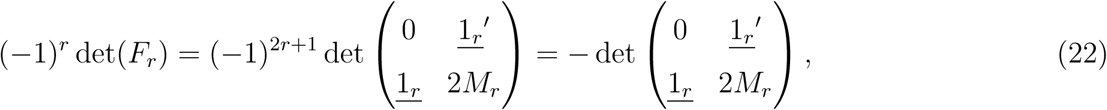

where *M*_*r*_ is the *r* × *r* matrix consisting of the upper-left corner of matrix *P*′*P*, and <u>1_r_</u> is the column vector of length *r* consisting of 1s.

We now apply a result for the determinant of partitioned matrices (Graybill, 1976, pp. 19-20). If *W* is invertible, then

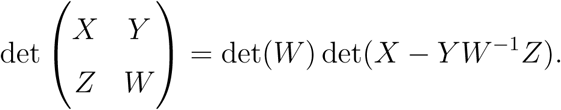

Applying this result to eq. 22, we obtain

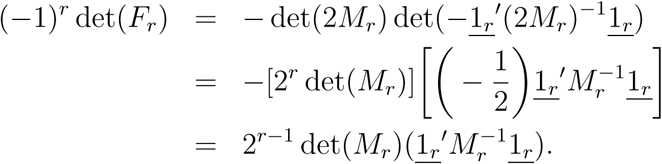

Because *P*′*P* is positive definite, *M*_*r*_ is also positive definite. To demonstrate this result, note that because *<u>x</u>*′*P*′*P<u>x</u> >* 0 for each nonzero column vector *<u>x</u>, <u>x</u>*′*P*′*P<u>x</u> >* 0 for each nonzero *<u>x</u>* with *x*_*k*_ = 0 for *k > r*. Because *M*_*r*_ is positive definite, det(*M*_*r*_) > 0 and 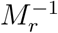 is also positive definite, leading to 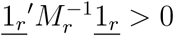. We conclude

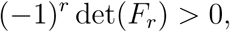

so that the critical point is the location of a local maximum.□

*Concluding the proof of Theorem 5.* Returning to part (iii) of the proof, following Lemma 10, if 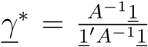 is interior to the simplex Δ^*K*−1^, then *H*_adm_ is maximal at *<u>γ</u>* = *<u>γ</u>*\*, with maximum 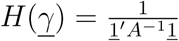. This value is the reciprocal of the sum of the elements of *A*^−1^. If *<u>γ</u>*\* is not interior to Δ^*K*−1^, then the maximum lies on the boundary of Δ^*K*−1^.

Finally, we note that 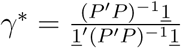 by using eq. 20. □

*Proof of Corollary 6.* In Theorem 5, the maximum of *H*_adm_ occurs either in the interior of the simplex Δ^*K*−1^ or on its boundary, {*<u>γ</u>* : <u>1</u>′*<u>γ</u>* = 1 and *<u>γ</u> ∈* Δ^*K*−1^}.

The boundary of the simplex is the union of *K* faces, which are themselves (*K* − 2)- simplices. If the maximum lies on the boundary of Δ^*K*−1^, then without loss of generality, we can permute the labels of the source populations so that *γ*_*K*_ = 0.

We drop column *K* from matrix *P* and apply Theorem 5 with this new *J* × (*K* − 1) matrix, *P*_{1,…,*K*−1}_, which has rank *K* − 1. By assumption, <u>1</u>′(*P*′{1,…,*K*−1}*P*{1,…,*K*−1})^−1^<u>1</u> ≠1.

We then apply Theorem 5 to *P*_{1,…,*K*−1}_. The maximum of *H*_adm_ occurs either at the point *γ*_*S*_, where *𝒮* = {1, 2, …, *K* − 1}, or on the boundary of the set {*<u>γ</u>* : <u>1</u>′*<u>γ</u>* = 1 and *<u>γ</u> ∈* Δ^*K*−2^}. We repeat this method of descent, decrementing the dimension (and permuting population labels without loss of generality) until we reach the case of only two source populations. A final application of Theorem 5 then finds that *H*_adm_ is maximized either interior to the 1-simplex—the line connecting vertices (1, 0) and (0, 1)—or at one of these vertices. □

## Appendix 2. Proofs for *K* = 2: Proposition 7 and Corollary 8

*Proof of Proposition 7.* We maximize the quadratic polynomial in eqs. 12-14 over *γ ∈* [0, 1]. The maximum occurs at the unique critical point or on the boundary of the interval.

Setting the derivative of eq. 14 with respect to *γ*_1_ to 0, we find that the critical point is

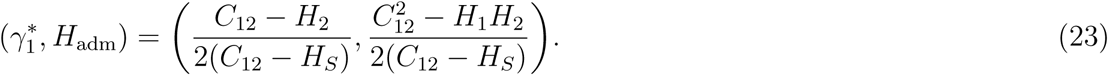

Because the leading coefficient of eq. 14 is negative for *<u>p</u>*<u>_1_</u> ≠ *<u>p</u>*<u>_2_</u>, the critical point is a maximum. Hence, if (*C*_12_ − *H*_2_)/[2(*C*_12_ − *H*_*S*_)] ∈ (0, 1), then the maximum of *H*_adm_ on the interval [0, 1] lies at *γ*_1_ = (*C*_12_ − *H*_2_)/[2(*C*_12_ − *H*_*S*_)]. Otherwise, the maximum lies either at *γ*_1_ = 0, in which case it equals *H*_2_, or at *γ*_1_ = 1, in which case it equals *H*_1_.

The conditions describing the location of the maximum can be written in terms of *H*_1_, *H*_2_, and *C*_12_. Because the denominator of 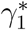 in eq. 23 is always positive for *<u>p</u>*<u>_1_</u> ≠ *<u>p</u>*<u>_2_</u> (Section 4), 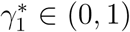 becomes equivalent to *C*_12_ > *H*_1_ and *C*_12_ > *H*_2_, the former inequality arising from the condition 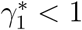 and the latter from the condition 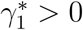.

If the requirement *C*_12_ > *H*_1_ and *C*_12_ > *H*_2_ for 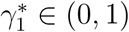 fails, then the maximum occurs on the boundary of the unit interval. We have *H*_adm_(0) = *H*_2_ and *H*_adm_(1) = *H*_1_. Thus, the maximum lies at *γ*_1_ = 0 if *H*_2_ > *H*_1_ and at *γ*_1_ = 1 if *H*_1_ > *H*_2_.

If *C*_12_ > *H*_1_ and *C*_12_ > *H*_2_ do not both hold, then one of them must hold, as we showed in Section 4 that 2*C*_12_ > *H*_1_ + *H*_2_. Combining the fact that either *C*_12_ > *H*_1_ or *C*_12_ > *H*_2_ holds with the observation that *H*_2_ > *H*_1_ leads to a maximum at *γ*_1_ = 0 and *H*_1_ > *H*_2_ leads to a maximum at *γ*_1_ = 1, we complete the characterization of the three cases. □

Note that alternative expressions in terms of *H*_1_, *H*_2_, and *F*_12_ can be derived by noting that 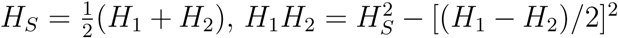 and *C*_12_ = *H*_*S*_(1 + *F*_12_)/(1 − *F*_12_), the latter relationship simply restating eq. 4 (recalling *C*_12_ = 1 for *F*_12_ = 1). Thus, we have

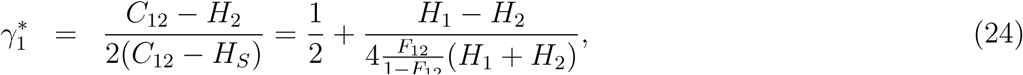

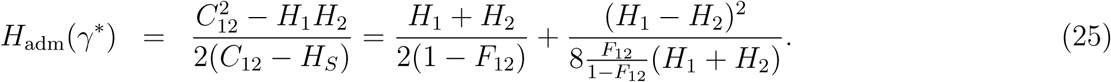

Another formulation uses the heterozygosity of a population formed by equal admixture of populations 1 and 2, or *H*_*T*_. Because *F*_12_ = 1 − *H*_*S*_*/H*_*T*_ by eq. 1, *F*_12_/(1 − *F*_12_) = (*H*_*T*_ − *H*_*S*_)*/H*_*S*_. Using this relationship in eqs. 24 and 25:

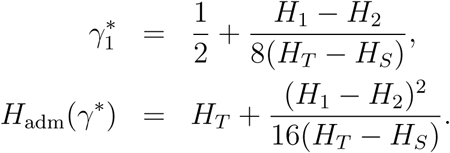

*Proof of Corollary 8.* We restate the condition 0 *<* (*C*_12_ − *H*_2_)/[2(*C*_12_ − *H*_*S*_)] *<* 1 as

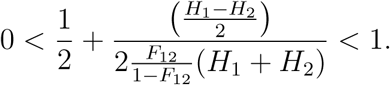

Subtracting 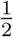 from both sides and multiplying by 2, an equivalent condition is

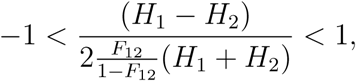

or, equivalently, 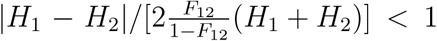. We rearrange this last expression to obtain the desired result. □

## Appendix 3: Dirichlet model for allele frequencies

We first provide results concerning *H*_adm_ in the case that the *K* source populations have independently and identically distributed (IID) allele frequency vectors. Next, we specify these IID vectors to be Dirichlet distributions.

### IID allele frequency vectors

We begin by examining the expected values of *H*_*k*_ and *H*_adm_.

#### Proposition 11.

Suppose the allele frequency vectors *<u>p</u>*_*<u>k</u>*_ are independently and identically distributed for 1 ⩽ *k* ⩽ *K*. Then 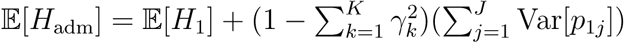.

*Proof.* We use eq. 8:

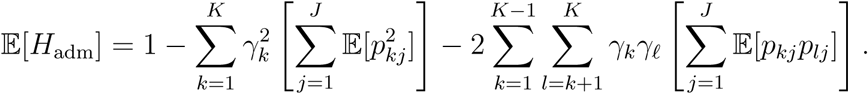

Using the IID assumption and simplifying by noting that 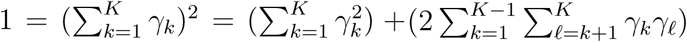, we have

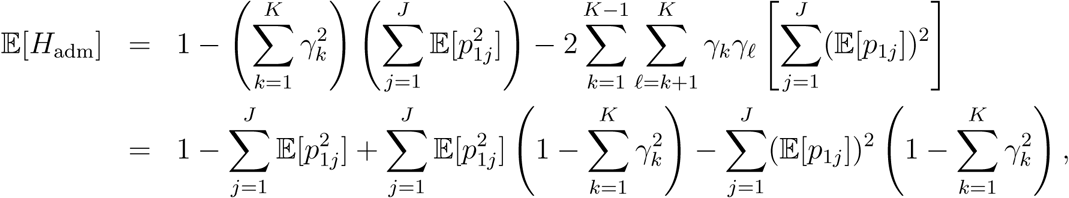

from which the result follows.

An immediate corollary of Proposition 11 is that under the IID assumption, *H*_adm_ has expectation greater than or equal to the expectation of the heterozygosity of each of the source populations.

#### Corollary 12.

Suppose the allele frequency vectors *<u>p</u>*_*<u>k</u>*_ are independently and identically distributed for 1 ⩽ *k* ⩽ *K*. Then 𝔼 [*H*_adm_] ⩾ 𝔼 [*H*_*k*_].

A second corollary results from the Cauchy-Schwarz inequality, by which 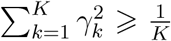, with equality if and only if 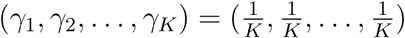.

#### Corollary 13.

Suppose the allele frequency vectors *<u>p</u>*_*<u>k</u>*_ are independently and identically distributed for 1 ⩽ *k* ⩽ *K*. Considering all admixture vectors *<u>γ</u> ∈* Δ^*K*−1^, 𝔼 [*H*_adm_] is maximized at 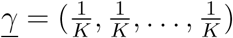, and has maximal value 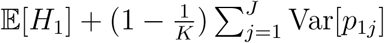.

### IID allele frequency vectors from a symmetric Dirichlet distribution

We now further assume that the independently and identically distributed allele frequency vectors follow a symmetric multivariate Dirichlet distribution. This distribution is frequently used for allele frequency distributions (Balding & Nichols, 1995; Pritchard *et al.*, 2000; Huelsenbeck & Andolfatto, 2007), and it is a natural probability distribution to assume for allelic types with the same marginal distributions.

The *J*-dimensional Dirichlet-(*α*_1_, *α*_2_, …, *α*_*J*_) distribution is defined over the open unit (*J* − 1)-simplex Δ^*J*−1^ and has concentration parameters *α*_*j*_ > 0. The means and variances for the individual allele frequencies are (Lange, 1997; Kotz *et al.*, 2000, chapter 49):

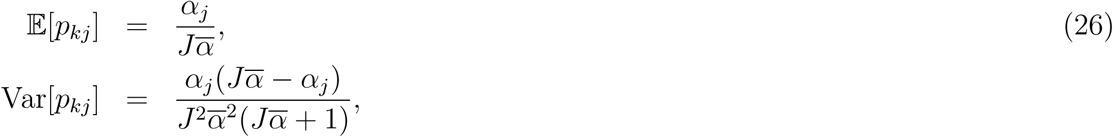

where 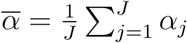.

The symmetric Dirichlet distribution assumes 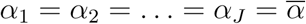, leading to:

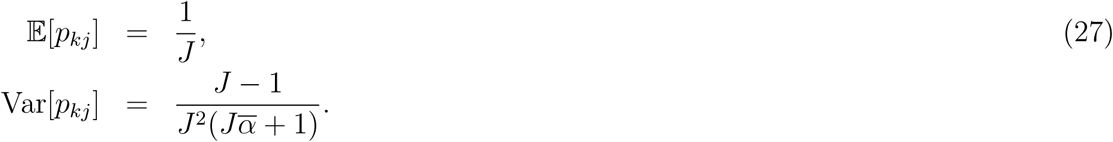

Making these substitutions in Proposition 11, we obtain the expectation of *H*_adm_ under the assumption that the allele frequency vectors follow independent Dirichlet distributions.

#### Corollary 14.

Suppose the allele frequency vectors *<u>p</u>*_*<u>k</u>*_ are independently and identically distributed for 1 ⩽ *k* ⩽ *K*, all with symmetric multivariate Dirichlet distributions with concentration parameter 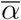. Then

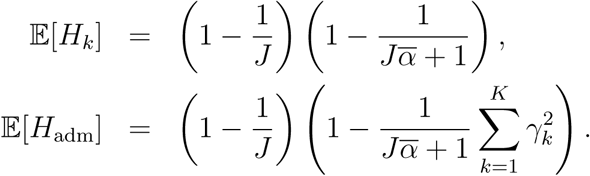

This corollary implies that both 𝔼 [*H*_*k*_] and 𝔼 [*H*_adm_] are increasing functions of *J* and 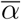.

The next proposition considers the special case of *K* = 2 and *J* = 2, further specifying a uniform distribution for *γ*_1_.

#### Proposition 15.

Consider *K* = 2 and *J* = 2. Suppose that the values of *p*_11_ and *p*_21_ are independently chosen from a uniform-[0,1] distribution. Suppose also that *γ*_1_ is also chosen from a uniform-[0,1] distribution. Then ℙ [*H*_adm_(*γ*_1_) > max{*H*_1_, *H*_2_}] = 1 − log 2 ≈ 0.307.

*Proof.* Using Proposition 9, we identify the regions of the unit square for (*p*_11_, *p*_21_) in which max_*γ* 1 ∈(0,1)_ *H*_adm_(*γ*_1_) > max{*H*_1_, *H*_2_}. These regions are 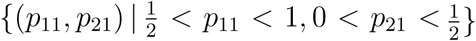 and 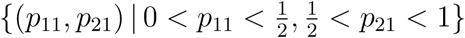.

Within those regions, we must determine the portion of the unit interval for *γ*_1_ in which *H*_adm_(*γ*_1_) > max{*H*_1_, *H*_2_}. *H*_adm_(*γ*_1_) is a quadratic function of *γ*_1_. We ignore the set of zero volume with *H*_1_ = *H*_2_. In the regions for (*p*_11_, *p*_21_) in which max_*γ*1∈(0,1)_ *H*_adm_(*γ*_1_) > max{*H*_1_, *H*_2_} and *H*_2_ > *H*_1_, the interval for *γ*_1_ in which *H*_adm_(*γ*_1_) > *H*_1_ is 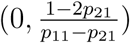. In the regions for (*p*_11_, *p*_21_) in which max_*γ*1_∈(0,1) *H*_adm_(*γ*_1_) > max{*H*_1_, *H*_2_} and *H*_1_ > *H*_2_, the interval for *γ*_1_ in which *H*_adm_(*γ*_1_) > *H*_1_ is 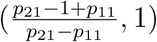.

The desired probability is the volume within the unit cube for (*p*_11_, *p*_21_, *γ*_1_) of the regions in which *H*_adm_(*γ*_1_) > max{*H*_1_, *H*_2_}. The volume is

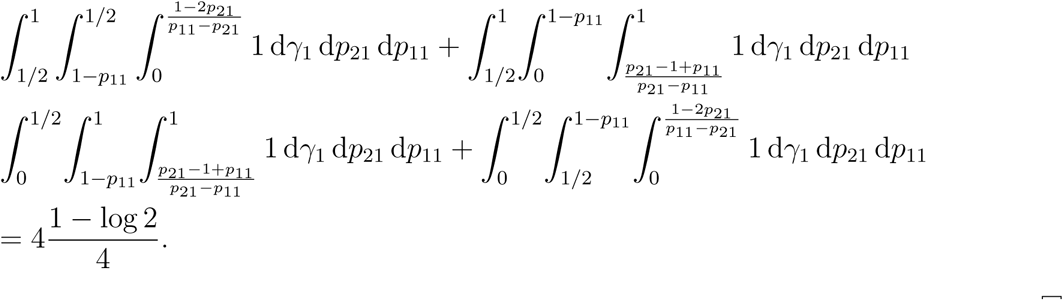

□

